# A LARGE-SCALE EVOLUTIONARY AND STRUCTURAL ANALYSIS OF CLC CHANNELS AND TRANSPORTERS

**DOI:** 10.1101/2025.07.07.663545

**Authors:** Ayush Mishra, Gladys Díaz Vázquez, Janice L. Robertson

## Abstract

The CLC family of membrane transport proteins consists of chloride channels and anion/proton antiporters that share a similar structural scaffold. How the same fold accommodates two fundamentally distinct mechanisms is poorly understood, and while the current set of experimental structures provide some information, the changes appear limited. In this study, we show that it is possible to scale up the structural information available using AlphaFold2 predictions and identify additional structural differences associated with each mechanistic class. A phylogenetic analysis is carried out across all known CLC genes to expand the classification to include 569 channel and 1,051 transporter homologs that have been modeled. Using distance matrices, we validate AlphaFold2’s ability to detect subtle structural differences among experimentally determined CLCs and use a random forest classifier to predict CLC channel vs. transporter sub-types to learn the structural changes that are the most important in the decision. The structural changes identified overlap with and contextualize the changes observed in experimental structures, expanding structural information across sequence space. The highest ranked change includes a contraction of distances between dimerization interface helices αH, αI, αP & αQ relative to the subunit core in the channel sub-types. This study lays out an approach for quantitative, large-scale structural analyses beyond experimental data and paves the way towards structural studies expanding on different conformational states and other protein families.

## INTRODUCTION

The CLC membrane transport protein family is nearly ubiquitous across biology, consisting of chloride ion channels and secondary active anion/proton antiporters. These membrane proteins play vital physiological roles governing excitability of muscle and neurons (Bennetts et al., 2005; Guzman et al., 2014), acidification of lysosomes and bones (Kornak et al., 2001; Zifarelli, 2022), and acid resistance in certain bacteria (Gut et al., 2006). Humans have 9 different versions of CLCs demonstrating these differential functions; mutations of these proteins lead to a variety of diseases, such as Bartter’s syndrome or muscle myotonia (Stölting et al., 2014). Even *E. coli* have three CLC genes: CLC-ec1, CLC-ec2, and the uncharacterized ecYfeO (Blattner et al., 1997; Iyer et al., 2002). CLCs therefore play important roles in human physiology but also across all domains of biology.

The last 20+ years have shed light on the structural architecture of CLCs. The first published structures were of CLC-st1 and CLC-ec1 (Dutzler et al., 2002) (**Fig. 1A**), containing 18 helices, αA-R, of which 16 are fully membrane embedded, αB-Q (**Fig. 1B**). Common to many transporters (Pornillos & Chang, 2006), CLC adopts an inverted-topology domain repeat with helices αB-I forming the first half, and αJ-Q forming the second (Forrest & Rudnick, 2009). The dimerization interface is formed by helices αH, αI, αP, & αQ (**Fig. 1C**) providing a thinned and curved hydrophobic surface that drives subunit association due to the free energy gained by burying this perturbative interface away from the lipid bilayer (Chadda et al., 2021). So far, all wild-type CLCs have been resolved as dimeric complexes like CLC-ec1. Dimerization is not required for function, as both monomerization and cross-linking the dimer interface still allows for Cl^-^ transport activity (Nguitragool & Miller, 2007; Robertson et al., 2010). In fact, each subunit contains its own permeation pathway that is formed along the symmetry axis of the inverted topology repeat, whereupon chloride ions are coordinated by several residues and supported by the electrostatic influence of the dipoles from helices αD, αF, αN, and αR. In CLC-ec1, residues S107 (helix αD) and Y445 (helix αR) coordinate chloride at the central binding site, with E148 (helix αF) serving as a gate for the external site (Dutzler et al., 2003).

**Figure 1.**
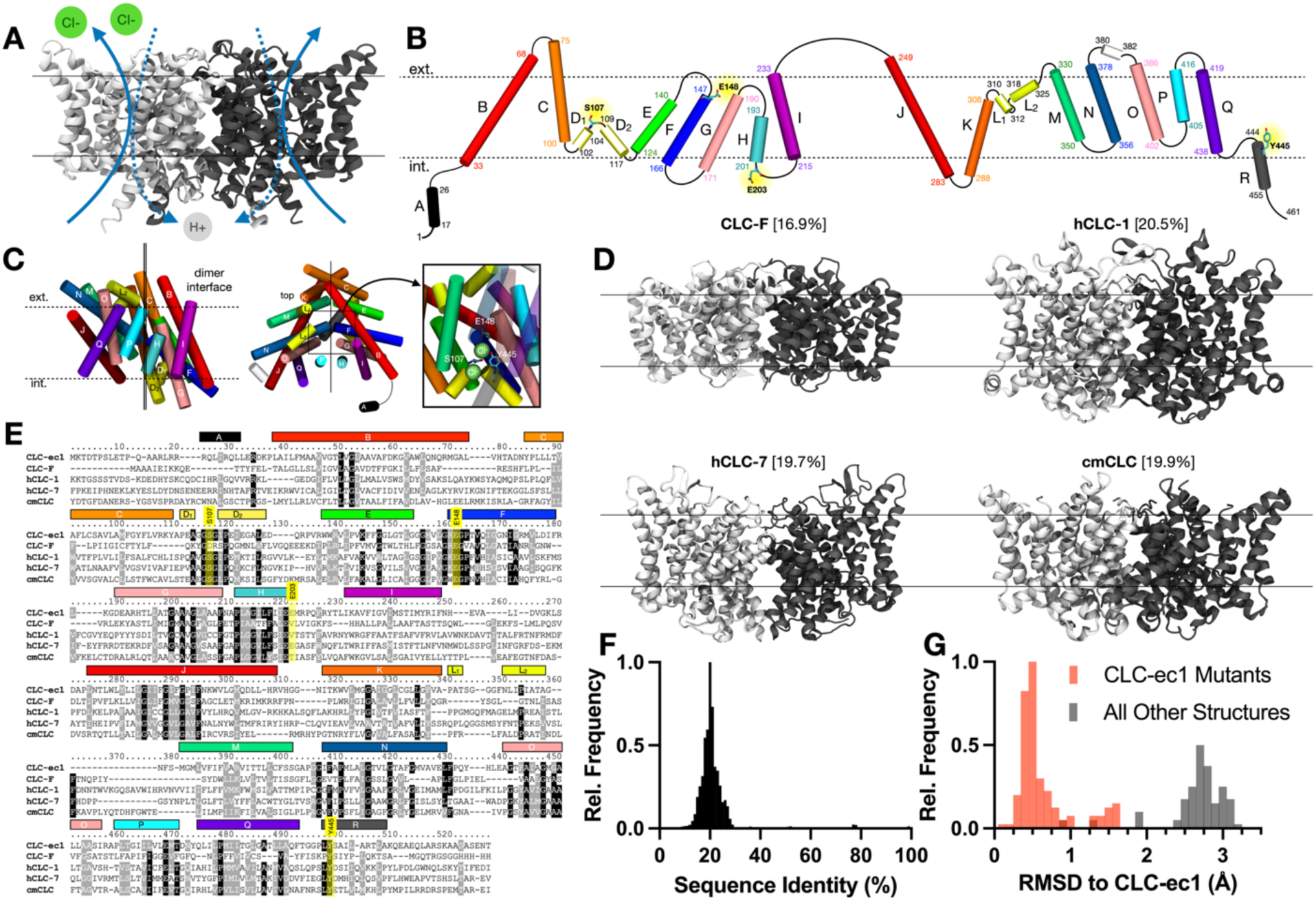
Sequence and structural diversity of CLCs. **(A)** CLC-ec1 dimer (PDB ID: 1OTS), with arrows illustrating 2:1 proton-coupled chloride transport occurring independently in each subunit. Approximate membrane interface positions are depicted by lines. **(B)** Topology of CLC-ec1 helices αA-R, maintaining the approximate orientation of each helix in the folded subunit, inside the membrane plane (dotted lines). Each helix is color-coded to highlight the inverted topology repeat. Numbers indicate flanking residue positions and key functional residues are highlighted and shown. **(C)** Orthogonal views showing transmembrane helix positions in the folded subunit. The axis of inverted symmetry is depicted as solid lines. *Left*, view of the dimer interface αQ, αP, αH & αI. *Center*, top view highlighting the region containing the transport pathway (box). *Right*, zoom in view of the transport pathway from inside the membrane plane, highlighting bound chloride ions and key residues: S107, Y445 and E148. **(D)** Structures of the membrane embedded regions of the same CLC homologs: CLC-F (PDB ID: 6D0J), hCLC-1 (PDB ID: 6COY), hCLC-7 (PDB ID: 8HVT), and cmCLC (PDB ID: 3ORG), with sequence identity compared to CLC-ec1 in brackets. **(E)** Multiple sequence alignment of representative CLC homologs highlighting shared sequence identity (dark grey) and similarity (light grey). **(F)** Histogram of the sequence identity of each CLC homolog from the InterPro database to CLC-ec1. **(G)** Root mean squared deviation (RMSD) of all C_α_ atoms of the solved structures compared to CLC-ec1 (PDB ID: 1OTS); mutant CLC-ec1 structures are represented in red.

When examining the greater CLC family, we see that it exhibits significant sequence diversity (**Fig. 1D**) despite sharing a common fold, inverted-topology architecture and dimer assembly (**Fig. 1E**). Indeed, homologous structures have sequence identity values as low as 15-25% when compared to CLC-ec1 (**Fig. 1F**). This occurs even for genes expressed within the same organism: *E. coli* contains CLC-ec1, CLC-ec2 and the uncharacterized ecYfeO. The sequence homology is low with CLC-ec2 sharing 21% sequence identity with CLC-ec1, and only 14% for ecYfeO. Examining the 14 unique CLC genes whose structures have been experimentally solved, we see that the fold is conserved along with the inverted topology repeat, while RMSD analysis of C_α_ backbone atoms shows 2.5-3 Å global deviations compared to CLC-ec1 indicating some conformational diversity (**Fig. 1G**). This is likely attributed to sequence variation, as similar analysis of CLC-ec1 mutant structures yields smaller changes 0.5-1.5 Å RMSD. This leads us to wonder if there are more structural changes observed within the CLC family that can be linked to the functional diversity.

CLCs are a family of membrane transport proteins that exist in two primary mechanistic forms: chloride channels and anion/proton antiporters. Ion channels and secondary active transporters involve distinctly different mechanisms in facilitating the movement of species across the membrane. Channels involve gating of an open, aqueous pathway that is accessible to both sides of the membrane such that ions diffuse passively down their concentration gradient. On the other hand, secondary active transporters require discriminatory access across the membrane, i.e., there is no continuous path for substrate diffusion at any point in the transport cycle. With this, the conformational changes required are linked to the binding of specific species, and with specific stoichiometry. The specific linkage between binding and conformational equilibria, enables the movement of one species down its concentration gradient coupled to the movement of another uphill, i.e., against its concentration gradient.

CLC-ec1, and all other prokaryotic CLCs whose structures have been solved have been demonstrated to function as secondary active transporters. However, eukaryotic CLCs exhibit both channel and transporter variants. Among human CLC homologs, hCLC-1, hCLC-2, hCLC-Ka, and hCLC-Kb have been shown to function as channels (Stölting et al., 2014), identified by accelerated rate of anion permeation and lack of proton transport linked to chloride permeation (Accardi & Picollo, 2010). The others, hCLC-3, hCLC-4, hCLC-6 and hCLC-7 have been demonstrated to act as coupled chloride/proton antiporters (Guzman et al., 2013; Leisle et al., 2011; Neagoe et al., 2010; Picollo & Pusch, 2005; Scheel et al., 2005). The structures of CLCs, whether they are channel or transporter type, have similar fold and topology. There are no obvious structural changes that indicate a different mechanism, as structures appear to be similarly dissimilar (**Fig. 1G**). Further, although certain residues are more common in each subtype, there are no clear sequence motifs that are predictive of channel or transporter activity (Fortea Verdejo, 2021). Sequence analysis shows that CLC channels often contain a valine at CLC-ec1 position 203, while transporters contain a glutamate (**Fig. 1E**). However, an ionizable side-chain is not required, as cmCLC contains a threonine at this position and still functions as a secondary active transporter (Feng et al., 2010a). Introducing a valine substitution at this position is also not sufficient to confer channel activity in CLC-ec1 (Lim & Miller, 2009), suggesting that other structural changes must be involved beyond this site. Indeed, mutating the external glutamate gate and central binding site residue tyrosine (E148 and Y445) to smaller side-chains creates a wide aqueous pathway across the membrane (Jayaram et al., 2008), increasing the rate of chloride transport and effectively creating a channelized transporter hybrid. While this shows that it is possible to introduce channel-like activity on a CLC structural template, these engineered substitutions are substantial, equivalent to boring a hole through the protein. They do not necessarily provide an understanding of the naturally occurring differences between CLC channels and transporters, as both residues are still well-conserved in the two subfamilies. Structural studies of CLC channels shed further light on this. In the CLC-K channel, a wider permeation pathway is observed along with flexibility of the dimer complex (Park et al., 2017), and in CLC-1, a unique conformation of the external site glutamate and widening of the permeation pathway are predicted to lower the kinetic barrier for ion permeation (Park & MacKinnon, 2018). It appears that there is a common theme of pore widening associated with channels, but that this may be conferred in different ways depending on the specific CLC sequence, making it challenging to identify based on structural data alone.

We hypothesize that examination of a larger number of CLC structures across sequence space will enable the identification of structural changes linked to CLC function that are not apparent with the available experimental structure data. To expand on the structural data set available, a phylogenetic analysis across all known eukaryotic CLCs was carried out to classify transporters and channels. Distance matrices (DMs) and distance difference matrices (DDMs) were used to validate the corresponding AlphaFold2 (AF2) structural models (Jumper et al., 2021) and their ability to detect structural changes between CLC homologs. Next, we used an ensemble-based machine learning approach to train random forest classifiers on an expanded set of 1,620 structural models of eukaryotic CLCs to quantify the importance of structural changes linked to CLC transporter or channel predictions. The results show that the most significant changes that are linked to changes in function involve reduction of distances between dimerization interface helices αH, αI & αP and the helix bundle αN, αO & αQ or αC in the subunit assembly. The structural changes observed in the AF2 models are also present in the experimental structural data set but are hard to identify as the most important changes do not always correspond to the largest structural changes. While this analysis likely focuses on CLCs in closed or occluded states, it shows that increasing the structural dataset with AF2 models can be used to identify subtle and distributed structural changes that are linked to differences in mechanistic function. This analysis can be expanded to examine other conformations through refinement of AF2 predictions or can be applied to any protein family of interest.

## RESULTS

### A phylogenetic analysis of eukaryotic CLC channels and transporters

At the onset of this study, there were nearly 75K CLC sequences identified in the InterPro CLC database (Blum et al., 2025) with only 13 genes represented by 106 experimentally solved structures, 55 of which were classified as wild-type (**Supp. Table 1**). The structures included 4 prokaryotic and 9 eukaryotic genes, and of the eukaryotic homologs, 6 genes are characterized as secondary-active transporters while 3 are ion channels. The general topology and homodimer assembly appear conserved amongst all CLCs (**Fig. 1D**), despite the low sequence homology (**Fig. 1E-F**). While RMSD analysis shows that there are global structural changes (**Fig. 1G**), we aimed to identify the specific changes in helix positions that are specific to CLC channel and transporter subtypes. However, the limited size of the experimental data set may obscure the identification of changes linked to mechanistic differences. Most of the InterPro sequences have low sequence homology compared to sequences with experimentally determined structures, around 20% (**Fig. 1F**), suggesting that there may be uncaptured structural variability in the family that can be gleaned from examining different sequences across evolutionary space. Recent advances in sequence-to-structure prediction using AF2 introduces a potential new source of information that can bridge this gap. Although AF2 tends to predict proteins in similar conformations, we hypothesize that the large number of predicted structures will expand structural variation over sequence space and will thereby increase the ability to identify structural changes linked to mechanistic differences in CLCs.

Before examining the AF2 models, we needed a way of genetically classifying sequences as CLC channels or transporters, as only a handful of CLCs have been experimentally characterized. To do this, we carried out a phylogenetic analysis to classify CLC sequences based on its evolutionary proximity to an already characterized sequence. We started with the ∼75K sequences in the InterPro database and filtered out highly similar sequences (95%+ sequence identity) and structural fragments, leaving 23,082 CLC sequences from all three domains of life (**Fig. 2A**). For the phylogenetic analysis, the dataset was further filtered for maximum 80% sequence identity, leaving 10,417 sequences. An approximate maximum likelihood tree was constructed with FastTree using a structural alignment to CLC-ec1 as the input multiple sequence alignment (**Fig. 2B**). There are two primary clades of eukaryotic CLC homologs: a large one that is close to CLC-ec1 and a smaller one near CLC-ec2. The larger clade contains all the eukaryotic homologs with known structure and likely evolved from an ancestral relative of CLC-ec1 in the last eukaryotic common ancestor. On the other hand, the smaller clade contains plant homologs like atCLC-e and atCLC-f which likely evolved from an ancestral relative of CLC-ec2. Beyond these two clades, there are a few other eukaryotic sequences scattered near ecYfeO, an uncharacterized CLC homolog also from *E. coli*. Further, archaeal sequences are scattered throughout the tree, a possible result of either very early horizontal gene transfer (Boto, 2009) or low phylogenetic signal.

**Figure 2.**
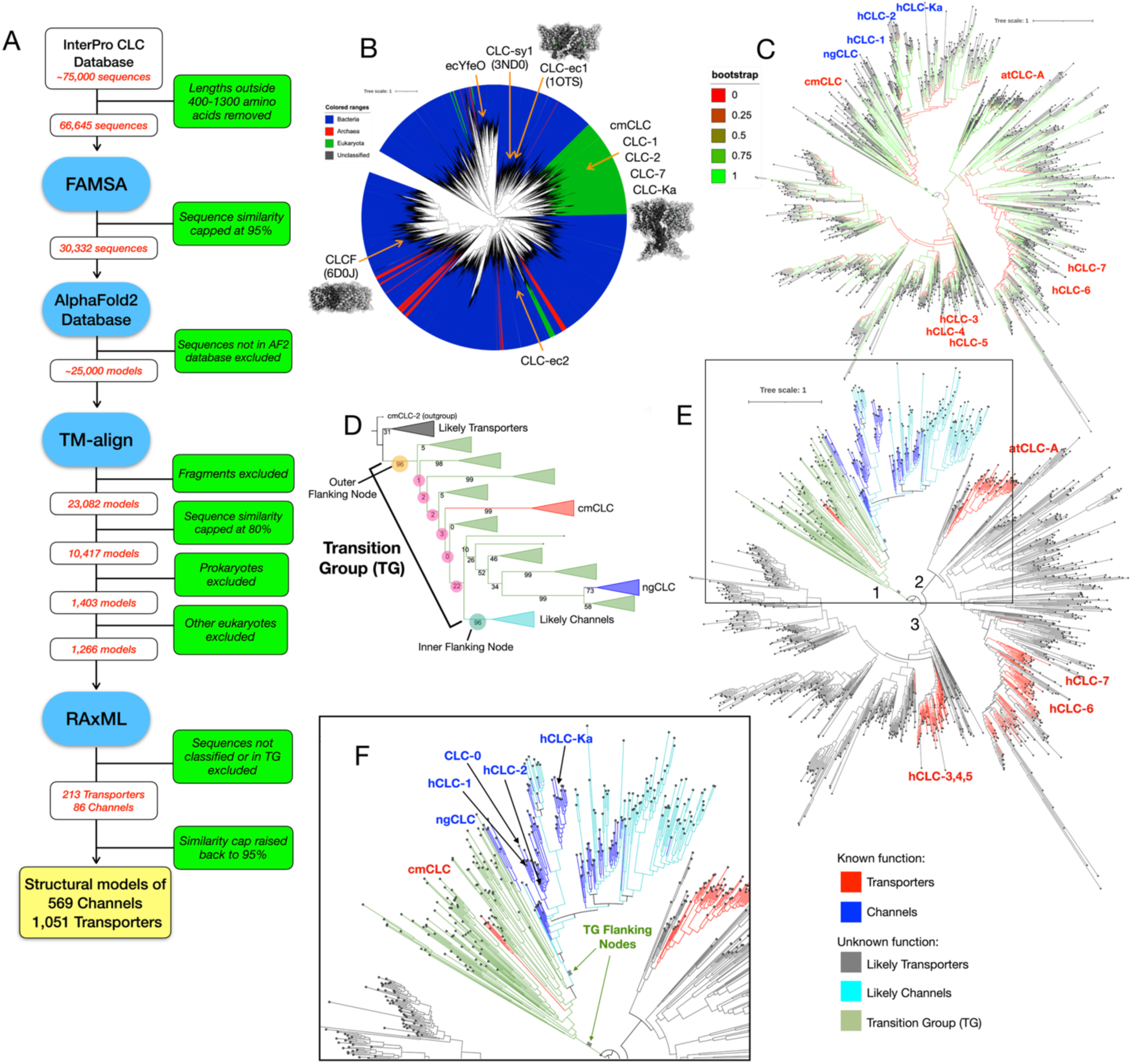
Phylogenetic analysis of the CLC Family. **(A)** Full sequence curation pipeline from IPR database to final model-fitting analysis. **(B)** Approximate Maximum Likelihood (ML) tree of 10,417 CLC sequences with FastTree; leaves colored by source organism domain and notable sequences are identified. **(C)** ML tree of 1,266 eukaryotic CLC sequences with RAxML from the main eukaryotic branch, showing bootstrap values. **(D)** Expansion of bootstrap support values along branch 1 showing a transition region that is demarcated by highly supported splits (both 96%) separating channels and transporters. **(E)** Sequences with known function and their proximate sequences are labeled according to channels, transporters or the transition group (TG). **(F)** Zoomed in view of branch 1 and transition group flanking nodes.

Since all known CLC channels are found exclusively within eukaryotic lineages, we extended our phylogenetic analysis to focus on the 1,266 sequences comprising the larger eukaryotic clade. A maximum likelihood tree, constructed using RAxML (Kozlov et al., 2019) and rooted with a sequence outside the clade, revealed three major branches (**Fig. 2C**). To assign functional categories, we mapped mechanistically characterized homologs onto the tree. Channel homologs include CLC-0 (Lísal & Maduke, 2008), hCLC-Ka/b (Kieferle et al., 1994), hCLC-1 (Saviane et al., 1999), hCLC-2 (Britton et al., 2005), and a putative channel ngCLC (Fortea Verdejo, 2021); transporter homologs include hCLC-3 (Guzman et al., 2013), hCLC-4, hCLC-5 (Picollo & Pusch, 2005; Scheel et al., 2005), hCLC-6 (Neagoe et al., 2010), hCLC-7 (Leisle et al., 2011), atCLC-a (De Angeli et al., 2006), and cmCLC (Feng et al., 2010b). The first branch contains all channel homologs and cmCLC, a known transporter, suggesting it also encompasses the transition from transporter to channel function. The second branch contains transporters: atCLC-a, hCLC-6 and hCLC-7. The final branch contains transporters hCLC-3, hCLC-4, and hCLC-5.

Based on the assumption that a functional switch did not occur recently in evolutionary history, we infer the function of uncharacterized sequences based on their evolutionary distance to these known homologs in both the final and bootstrap trees, resulting in 569 putative channels and 1,051 putative transporters.

Within the first branch, we identified a paraphyletic group, termed the *transition group* (TG) (Fortea Verdejo, 2021), flanked by two highly supported nodes (96% bootstrap support; **Fig. 2D**). This group includes both cmCLC (a transporter) and ngCLC (a putative channel), with poorly resolved internal structure indicating low phylogenetic signal. Because of the poor resolution, we group all 194 sequences within this region as transition group homologs. Further, assuming a single evolutionary transition from transporter to channel, we can infer that all sequences descending from the inner flanking node (toward ngCLC) are likely channels, whereas those outside the outer node (toward cmCLC) are likely transporters. We adopt this inference to create a second, expanded version of our functionally classified dataset to test the generalizability of our structural findings (**Fig. 2E, F**).

### Validation of AlphaFold2 CLC models for accuracy in structure and structural changes

With this expanded classification set, we examined AF2 predictions of CLC structures and their ability to predict structural changes between homologs by quantitative structural comparison. For this, we use distance matrices as a scalable way to compare structures of variable sequence and structure. We first calculate the distance matrix (DM) for each CLC, which provides a 2D quantitative map of the protein where each point on the matrix represents the cartesian distance between the Cα of each residue pair (**Fig. 3A**). While prior applications of DMs to the CLC family have largely focused on comparing alternative conformations of the same sequence (Chavan et al., 2020), their utility extends further. Because DMs are internally defined, they offer a robust measure of structural difference that is largely insensitive to the specific method of structural alignment, unlike traditional approaches like RMSD that require explicit structural superposition. To accommodate comparisons of different CLC homologs with varying sequence content, we focus our analysis on residues among transmembrane helix and other functional segments commonly found in most CLCs. This avoids loops, but includes residues in helices αB-αQ, the αC-D linker (i.e., residues 100-109 on CLC-ec1), and the N-terminal half of helix αR. A structural alignment of each CLC homolog was carried out with CLC-ec1 using TM-align and then alignment positions were removed that have less than 85% occupancy across the CLCs studied to generate a common list of *alignment sites* from 1-294. Since intra- and extracellular loops are highly variable, this focuses the analysis on slow-changing structural elements in the transmembrane regions resulting in the truncated and aligned distance matrix DM_aln_ (**Fig. 3B**).

**Figure 3.**
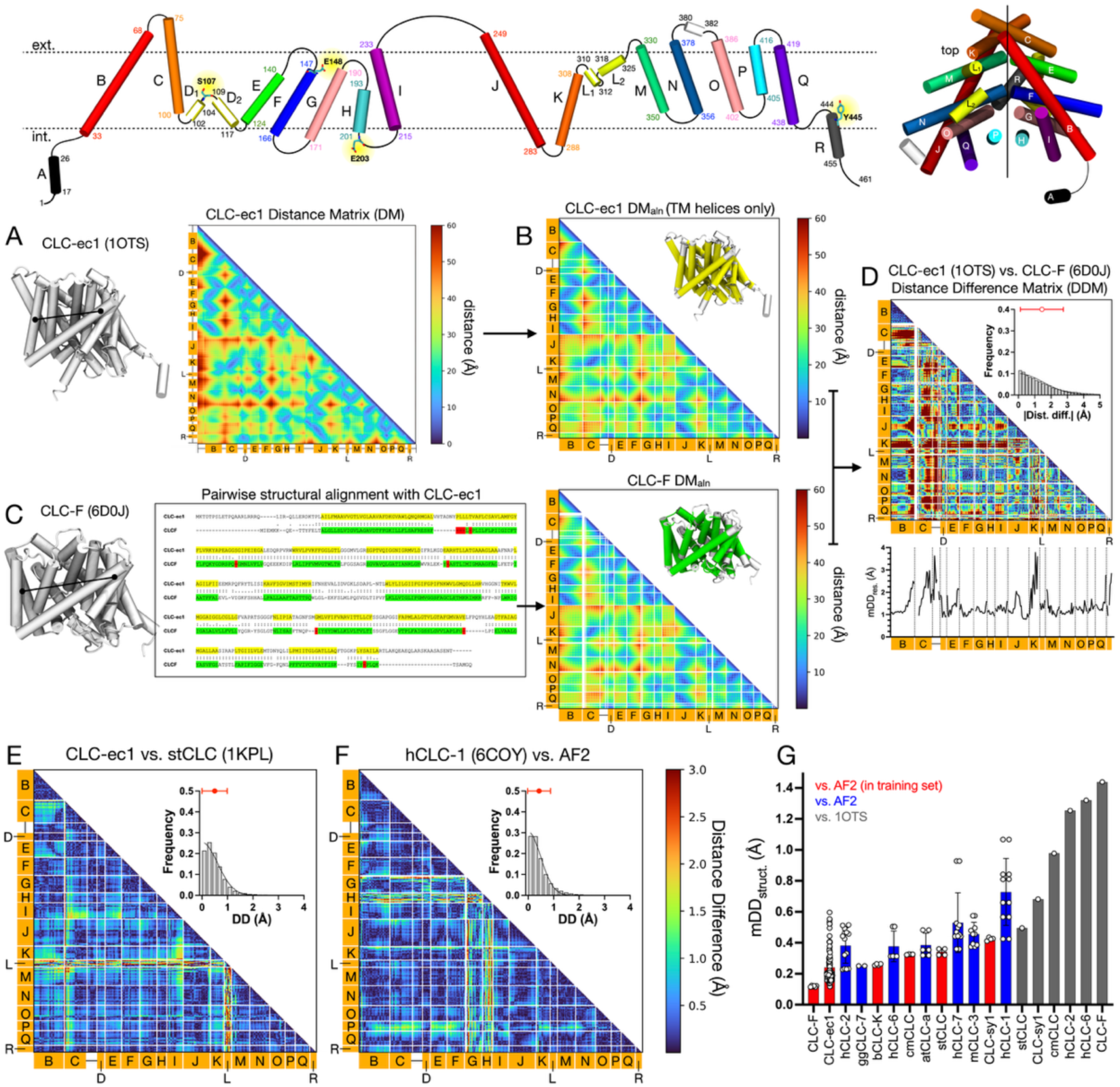
Quantifying structural differences between CLC structure using distance matrices. *Top*, the topology map and subunit assembly for CLC-ec1 is provided as a structural guide. **(A)** Residue-residue distance matrix (DM) for CLC-ec1 (PDB ID: 1OTS), computed from all pairwise C_α_ distances. **(B)** Aligned and truncated DMs featuring transmembrane helix segments along a common structural alignment, DM_aln_. White lines indicate the ends of transmembrane helices. **(C)** DM_aln_ for CLC-F (PDB ID: 6D0J), where filtered residues are annotated (green) via structural alignment with CLC-ec1; gaps in the structural alignment (red) create gaps in the resulting distance matrix (white lines). **(D)** Absolute distance difference matrix (DDM) between CLC-ec1 and CLC-F. Frequency histogram of absolute distance differences are shown in the inset plot, with mean DD_struct._ (mDD_struct._) ± std, a metric for comparing global distance differences, is plotted in red. Mean absolute distance differences by residue (mDD_res._) is shown as a line plot below. **(E)** DDM for stCLC (PDB ID: 1KPL) and CLC-ec1. **(F)** DDM for hCLC-1 (PDB ID: 6COY) and its AF2 predicted model. **(G)** Summary of mDD_struct._values ranked by minimum value, for experimental CLC structures compared to CLC-ec1 PDB ID 1OTS (grey), and CLC homologs compared to their AF2 predictions. The AF2 values are designated by whether the experimental CLC structure was eligible to be in the original training set (red) or solved after AF2 (blue).

The truncated DM_aln_ matrix can be used to identify structural differences between any two CLCs regardless of sequence homology. For example, to compare the fluoride/proton antiporter CLC-F with CLC-ec1 (17% shared sequence identity), we first generate a pairwise sequence alignment based on the structural alignment of CLC-F on CLC-ec1 and then map the CLC-F sequence onto the common *alignment sites* to determine the residues used in DM_aln_ (**Fig. 3C**). To compare between the two, we calculate the absolute value of the difference between two DM_aln_s to obtain an absolute distance difference matrix (DDM), where high values indicate structural changes between two aligned residue positions (Nishikawa et al., 1972). A DDM can be calculated between a CLC structure and its AF2 model, two different CLC homologs, e.g., the prokaryotic CLC-F vs. CLC-ec1 (**Fig. 3D, Supp. Fig. 1**), or two groups of CLCs, such as the differences between all known channels and transporters (**Supp. Fig. 4**). To summarize DDMs and identify regions that exhibit robust structural changes, the 2D matrix data is averaged along each alignment position to yield a line plot reporting the mean distance difference per alignment site, i.e., mDD_res._ (**Fig. 3D**). Furthermore, distance differences can be averaged across the entire matrix, mDD_struct._, to provide a global quantitative measure of the difference between two structures, analogous to an RMSD score in a traditional structure alignment. With this methodology in place, we can systematically quantify structural changes between CLCs that exhibit low sequence homology.

We calculated the DDMs between all experimental CLC structures compared to CLC-ec1 (**Fig. 3E; Supp. Fig. 1**) or their AF2 predicted model (**Fig. 3F; Supp. Fig. 2**). AF2 models exhibit minimum mDD_struct._ values between 0.1-0.4 Å for at least one of the experimental structures, indicating that AF2 is capable of successfully predicting a CLC structure for all homologs studied, including those in and out of the original training set (**Fig. 3G**). These differences are less than the value observed when comparing two experimental CLC structures that share high sequence identify, e.g., CLC-ec1 and stCLC at ∼80%, with mDD_struct._ = 0.5 Å (**Fig. 3E; Supp. Fig. 1**), indicating that AF2 predicts CLC structures with accuracy. This accuracy carries over to predicting distance difference changes between different CLC homologs (**Supp. Fig. 3A**), showing a correlation between structural changes predicted by AF2 and those observed in experiments. The correlation indicates that AF2 underpredicts these differences ∼ 2-fold (**Supp. Fig. 3B**), so changes identified as significant by AF2 are expected to also be observed in experimental structures. With this evidence of general success in predicting experimental CLC structures and structural differences, we rationalize the use of AF2 CLC models in our large-scale structural analysis.

### Classification of eukaryotic CLC channels and transporters by ensemble-based machine learning

The expanded set of CLC genes and structures associated with channel and transporter functions enables us to take an ensemble-based machine learning approach to identify features that are associated with mechanistic differences (**Fig. 4A**). Because machine learning models can capture complex, non-linear trends in the data, we trained a classifier to predict whether a CLC gene was a transporter or channel based on its structural model. For each of the 569 CLC channels and 1,051 CLC transporters, we compute the aligned distance matrix DM_aln_ and flatten them into one-dimensional vectors of 43,071 unique pairwise distances that serve as a machine-readable representation of a structure’s features. These vectors formed the training input for a Random Forest Classifier (RFC), a supervised machine learning algorithm that builds an ensemble of decision trees to assign class probabilities. In this case, the classifier is trained to predict the likelihood a given structure is a channel and outputs the probability *P_Chan._* The successful ability of the RFC to predict channels and transporters is based on its learning of structural differences. These learned differences can be quantified as the importance the classifier prescribes to each input feature. Consequently, these features highlight regions of the protein that are associated with the changes in mechanistic function.

**Figure 4.**
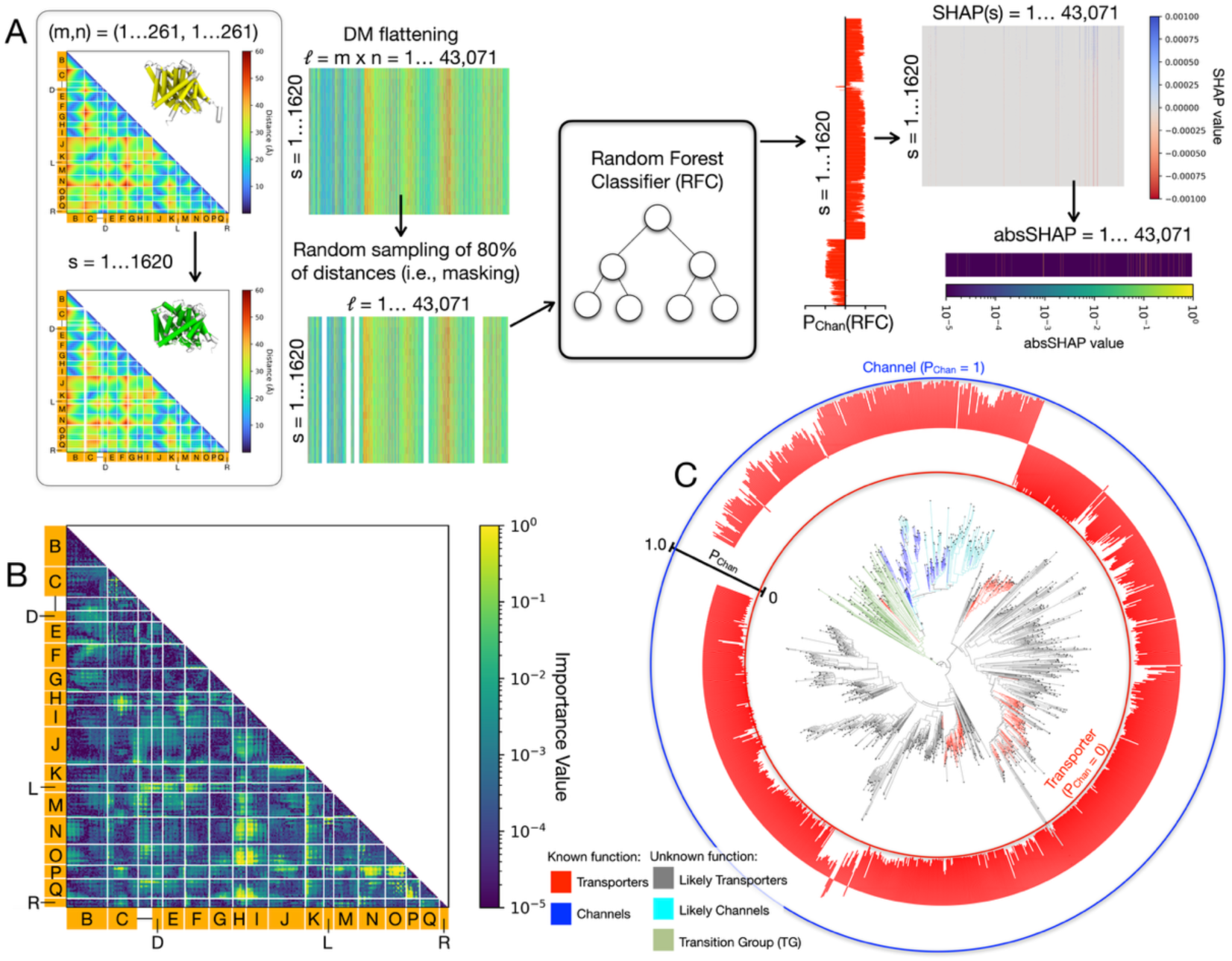
Structure analysis via large-scale model-fitting. **(A)** A graphic describing the model-fitting approach for large-scale structure analysis. **(B)** 2D map of importance values corresponding to the structural elements of CLCs. (**C**) Channel probabilities assigned by a Random Forest Classifier trained on likely transporters and channels without masking among CLCs in the eukaryotic phylogeny.

To interpret how a classifier makes its predictions, we employed SHAP (SHapley Additive exPlanations) values, a method that quantifies the expected marginal contribution of each input feature to the classifier’s output for a single prediction (Winter, 2002) (S. M. Lundberg & Lee, 2017). Absolute SHAP (absSHAP) values report the average contribution of each input feature across all structures used in the training data. To appropriately sample absSHAP values for all features, we trained 1,000 separate RFCs, each using a different random selection of 80% of the input features and define the *importance value* as the average of the absSHAP values over the 1000 RFCs for each input feature (**Fig. 4B**). These values represent how important each corresponding element of a structure’s distance matrix is to differentiating transporters and channels.

To evaluate the performance of the RFC in distinguishing CLC channels from transporters, we employed two complementary validation strategies. First, we performed 5-fold cross-validation on the training set to test for overfitting. The training dataset was partitioned into five equal subsets, and the classifier was trained on four subsets while the remaining subset was used for testing. This process was repeated for all partitions, providing a robust measure of classifier accuracy (Bates et al., 2024). In all cases, the unidentified sequences in the subset were appropriately classified, yielding a score of 1.0, indicating 100% accuracy on the ability of the classifier to interpolate sequence placement.

In a second validation test, we sought to determine how generalizable the identified structural trends were. We applied a single, fully trained RFC, i.e., using all features of all available training data, to predict channel probabilities across the broader eukaryotic phylogenetic tree (**Fig. 4C**). Assuming a single functional transition occurred from transporter to channel during eukaryotic CLC evolution, we expected the classifier to reflect three distinct trends across phylogenetic space. First, sequences outside of the outer flanking node of the transition group should be predicted as transporters, reflecting their ancestral state. Indeed, most of these sequences received channel probabilities close to zero, with a few exceptions where intermediate probabilities (∼0.5) were observed. Some examples include *Candida inconspicua* CLC (Uniprot ID: A0A4T0X126) and *Wickerhamiella sorbophila* CLC (Uniprot ID: A0A2T0FE06), both near hCLC-3; *Galdieria sulphuraria* CLC (Uniprot ID: M2WY17) near hCLC-7; and *Micromonas pusilla* CLC (Uniprot ID: A0A7S0D4I2) near atCLC-a. Such cases may reflect structural divergence from canonical transporter profiles, potentially indicating alternative or hybrid mechanisms. Second, sequences to the right of the inner flanking node of the transition group were expected to be classified as channels, reflecting their derived function. Here, the classifier performed as anticipated, assigning channel probabilities in the 0.9 to 1.0 range. Third, sequences within the transition group were expected to show intermediate behavior, consistent with a mixture or gradation of functional features. Indeed, these sequences exhibited a broad range of predicted channel probabilities, spanning from ∼0.5-0.8 with a slight upward trend as sequences got closer to the channel clade. This variability supports the hypothesis that these structures may represent intermediates or functionally ambiguous forms not fully captured by either class. Overall, the classifier met all three evolutionary expectations, indicating that it successfully generalized beyond the labeled training set and meaningfully extrapolated structural-function relationships across the phylogeny.

### A large-scale structural analysis of eukaryotic CLC channels and transporters

A robust classifier allows us to quantify importance values associated with each pairwise distance. By ranking the importance values, we can identify the most significant structural changes that factored into the classifier’s learning that a CLC structure corresponds to channel or transporter type function (**Fig. 4B**). Examining the top 1% of importance values (**Fig. 5A**) and clustering them according to helix positions allows for a ranking of 29 structural regions strongly associated with CLC channel or transporter sub-types (**Fig. 5B**). Seventeen out of the total 29 regions examined, including all in the top 10, include a helix that is located at the dimerization interface—αH, αI, αP or αQ. Comparing the distance distributions of these regions shows there are distinct structural changes between channel, transporter and transition group sequences (**Fig. 5C**). In channels, the most significant change involves a reduction in the distance of dimerization interface helices αH, αI & αP with the adjacent helix bundle αN, αO & αQ. Within this bundle, dimer interface helix αQ moves away from αN & αO in a small but significant change. Helix αK located on the opposite side of the subunit exhibits larger distances from the αN, αO & αQ bundle (now including αJ) due to a reduction of secondary structure in the intracellular region of αK in channel sub-types. Note, αK and αC form a helix-helix bundle stabilizing the two halves of the inverted topology assembly. The dimerization interface helices αH, αI & αP also move closer to αC reflecting a compression of the entire subunit assembly in channels, although this is not as well reflected among the available experimental structures. Finally, in channels, the distance increases between αL and αD & αF, i.e., around the external gate E148 in CLC-ec1, as well as αL and the intracellular part of αN. This reflects an expansion of the subunit in the direction along the membrane normal vector, near the chloride binding sites along the transport pathway. These structural changes are recapitulated in the experimental data set, with distance changes that overlap with the AF2 distributions (**Fig. 5C**) and are evident in structural alignments of the channel hCLC-1 (PDB ID: 6COY) and transporter hCLC-7 (PDB ID: 8HVT).

**Figure 5.**
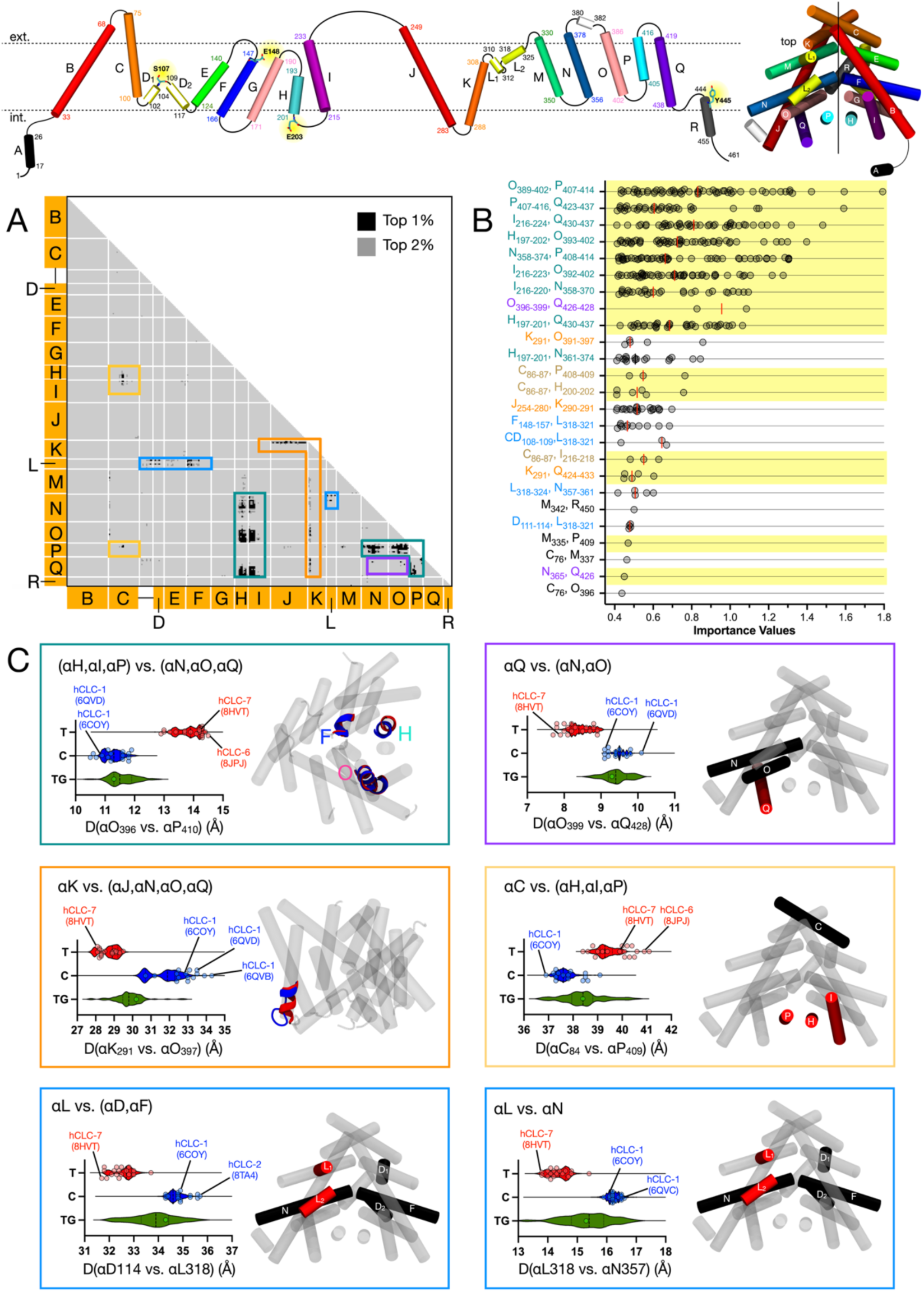
Structural differences between CLC channels and transporters associated with highest-ranked importance. *Top*, a cartoon showing CLC-ec1’s helical topology and subunit assembly as a guide. **(A)** Distance positions associated with the top 1% (black) & 2% (grey) importance values from the classification. Boxes indicate structural clusters. **(B)** Ranking of peak importance values grouped into structural clusters (median – red line). Regions containing at least one of the dimerization interface helices (αH, αI, αP or αQ) are highlighted yellow. **(C)** Largest importance value distance distributions for residue pairs in each cluster according to phylogenetic classification (Transporter (T, red), Channel (C, blue), Transition group (TG, green)). Distances from experimental structures are shown as overlaying scatter plots. Structural changes are shown for the transporter hCLC-7 (PDB ID: 8HVT) and channel hCLC-1 (PDB ID: 6COY) on the right.

To visualize these structural changes on the AF2 dataset, we selected a channel-transporter pair with the lowest number of structural changes in distances outside of the top 10% most important distances. *Betta splendens* bsCLC was selected as the channel candidate (Uniprot ID: A0A6P7PB93) and cfCLC from *Cavenderia fasciculata* as the transporter candidate (Uniprot ID: F4PTB4), whose average distance difference for the top 5% in importance is 1.57 Å and average distance difference for distances *outside* the top 10% is 0.49 Å (**Supp. Fig. 5A**). Morphing between these two representative structures illustrates the structural changes that are most important in determining channel or transporter mechanistic types (**Supp. Fig. 5B, Supp. Video 1**).

While this analysis picks up on the structural features that are most important for discriminating between channels and transporters, these are not necessarily the *largest* distance changes between structures. To determine these differences, we calculated a DDM between all AF2 models of channels and transporters used in the importance analysis (**Supp. Fig. 4C**) and found the top 1% of distance differences (**Supp. Fig. 4F**). The largest structural changes involve αK around residues 290/291 with a cluster of helices in the second half of the protein, αJ, αO, & αQ; this is the same cluster we observe to have relatively high importance. The next largest change involves the intracellular end of αJ bending towards the center of the structure, shortening the distance to helix αC. This change is even more pronounced in the larger set of AF2 models, indicating that the experimental dataset underrepresents this feature. Despite this, it does not fall into the top 1% of ranked importance values (Peak Importance Value = 0.26). Third, αH is observed to move inwards into the structure. Next, residue 148 in αF (i.e., the external glutamate gate in CLC-ec1) rotates upwards toward residue 319 on αL and away from residue 200 on αH. While the vertical movement of residue 148 towards αL has a high-ranking importance value, the movement away from αH is less so. Fifth, the intracellular region of αF shows a reduction in helicity in channels leading to large distance changes relative to a few residues, most apparent when measured from the extracellular end of αE. This feature is not picked up by the importance analysis, and the magnitude of this change is reduced when more sequences are used, indicating that the experimental data overrepresents this. While these reflect the largest magnitude changes, they are not all considered to be the most important features in predicting functional differences, typically because the distance distributions overlap (**Supp. Fig. 4J**).

## DISCUSSION

This study demonstrates that AF2 models can be used to expand on the structural information available, and identify features linked to mechanistic differences across the family of CLC channels and transporters. Although AF2 predictions are not without error and can deviate from experimentally determined structures in key regions, our findings suggest that these limitations are outweighed by the statistical power conferred by massively increasing structural sampling. By integrating hundreds of predicted models across diverse homologs, we can identify recurring, statistically robust features that correlate with functional divergence. CLCs are a particularly well-studied protein family, with about 75,000 known sequences and 106 total experimental structures available to date. Despite this, AF2 can corroborate major structural changes observed in the experimental data set and provide unique insights that are not readily apparent from the experimental structures alone. For other, less-studied protein families with few or no solved structures, the mechanistic and functional insights enabled by AF2 are likely to be even more impactful. Here, studying structure across these families allows us to bypass some of the limitations of sequence-based analyses and instead uncover conserved architectural features, functional motifs, and conformational signatures that may be obscured by billions of years of sequence variability alone.

The validation analysis shows that AF2 captures many of the same structural differences between CLC subtypes that are observed in experimental data, including shifts in dimer interface helices and changes around the permeation pathway. This indicates that while the method may not always produce perfectly accurate individual structures, it does preserve meaningful structural trends across billions of years of evolution. In this sense, AF2 enables a new kind of structural analysis: one that treats the problem statistically, using structural ensembles to extract mechanistic signals above the noise. Moreover, comparison of DM_aln_s serves as a structural analog to sequence-based evolutionary studies, where amino acid divergence is assessed through multiple sequence alignments. Therefore, this structural framework is not restricted to identifying differences between functional subfamilies but can be generalized to many of the same evolutionary inferences traditionally drawn from sequences alone. For instance, by examining how conserved a given structural region is across a diverse set of models, one can identify functionally important or conformationally constrained regions of the protein, analogous to conserved motifs in sequence alignments. Additionally, this approach enables the detection of regions that evolve more rapidly in specific clades, potentially highlighting areas under differential selective pressure or involved in lineage-specific adaptations. In this way, DM_aln_-based analyses can provide both structural insight and evolutionary context, offering a powerful complement to traditional sequence-based methods.

Our analysis does not explain all the structural distinctions that are meaningful between CLC channels and transporters. Rather, this large-scale analysis sheds light on structural commonalities that are evolutionarily conserved, keeping in mind that they may be correlative rather than causal. Surprisingly, we find the changes that are most important in factoring into the classifier’s ability to predict that a CLC is a channel involves the dimerization interface, a region that was hinted at in the experimental analysis but not overwhelmingly apparent. The positioning of this interface relative to the rest of the subunit, either helix bundle αN, αO & αQ or αC, indicates a global compression of the subunit in channels compared to transporters. Keeping in mind that the CLC structures in the original training set were likely partially closed channels or occluded transporters (Khantwal et al., 2016; Park et al., 2017), these results may inform on channels evolving a more compact fold in these states to prevent ion leakage. On the other hand, CLC transporters participating in alternate access conferred by side-chain rotamer changes may not require the same level of structural compression to occlude the pore. Indeed, structural changes in the dimerization complex were observed in CLC-K (Park et al., 2017), along with the positioning changes of αH in CLC-1 (Park & MacKinnon, 2018). While further structural analysis along the pore is warranted for a better mechanistic understanding, this would require examining the conformations of CLC channels and transporters that correspond to active states.

This highlights that a major limitation of our analysis is the inherent biasing of AF2 to predict a single, best-guess of the resulting structure. In this respect, we interpret our observations as associations informing on underlying changes in subunit fold, rather than the mechanistic states that lead to the physically different functions. The structures predicted by AF2 are biased to states represented in the PDB in the original training set, and experimental PDBs tend to be biased by the structures that are at the energetic minimum favoring certain states. Indeed, AF2 has been shown to be less effective at predicting the structure of fold-switching proteins (Chakravarty & Porter, 2022), revealing its inherent bias towards structures in the training set. Considering CLCs, prior to 2018, there were 6 CLCs in the PDB, and only one eukaryotic CLC that was classified as a transporter (cmCLC). Due to the nature of CLC-ec1 and other prokaryotic homologs lacking an aqueous pathway across the membrane, most of the conformations in the training set are expected to correspond to some type of occluded state (Khantwal et al., 2016). Even in CLC-K, there is a constriction along the aqueous pathway indicating this is likely a closed or inactive state of the channel (Park et al., 2017). As a result, structural differences associated with dynamic transitions, such as those observed at the αC-D linker, are clearly present in the experimental data set, but are largely absent in AF2 models and cannot be picked up in our analysis. However, this limitation seems to be addressable. Recent studies show that AF2 can be guided to sample alternative conformations through reducing MSA depth (del Alamo et al., 2022) or directly changing residues at certain alignment sites of the MSA (Stein & Mchaourab, 2022). Therefore, future applications of this approach could be extended to deliberately explore different conformational states, enabling a comparative analysis of state-dependent structural changes across homologs in a similar manner done in this study. Such efforts could offer even deeper insight into the mechanisms of channel vs. transporter activity within the CLC family, and more broadly, into the conformational dynamics underlying function in other systems.

Another limitation is that the power of this analysis hinges on the strength of the functional classification by phylogenetic analysis. Only a few structures have been experimentally characterized, typically assessed by electrophysiological approaches that measure the reversal potential of conductance/transport. Expansion of this functional classification required making a few assumptions. For example, during the validation of the random forest classifier results (**Fig. 4**) we assumed that there was only one functional transition from transporter to channel in the eukaryotic phylogeny. However, it is possible that other, more recent functional switches occurred. Such misclassifications would propagate through the model, potentially obscuring true structure-function relationships or introducing spurious ones. Future studies incorporating more extensive experimental validation will be critical to refining these assumptions and increasing the validity of large-scale functional predictions.

## ACKNOWLEDGEMENTS

The Robertson lab is supported by the National Institute of General Medical Science, National Institutes of Health (R01GM120260, R03NS133680). We thank Dr. Michael Landis and the members of the Robertson laboratory for helpful feedback during the development of this project.

## METHODS

### Large-scale CLC sequence curation

To efficiently analyze a large set of CLC sequences, the following sequence curation protocol was carried out to remove redundancies and allow for the analysis of many AF2 models together (**Fig. 2A**):

1. CLC sequences were retrieved from the InterPro database (IPR ID: IPR001807) containing ∼75,000 sequences at the onset of this study.
2. CLCs with lengths between 400-1300 residues were considered likely to be homologous to the CLC-ec1 membrane embedded domain, resulting in 66,645 sequences.
3. This set of CLCs were aligned with the *Fast and Accurate Multiple Sequence Alignment* (FAMSA) method and clustered at 95% sequence identity, retaining one representative per cluster. The representative was chosen to be either a predefined reference sequence or the sequence with the longest aligned length to CLC-ec1’s TM residues. This resulted in 30,332 remaining sequences.
4. In the next step, ∼5,000 sequences that were not in the AF2 database were removed, leaving ∼25,000 sequences.
5. All remaining structures were aligned to the AF2 model of CLC-ec1 using TM-align. Sequences with fewer than 94% occupied alignment sites among transmembrane residues of CLC-ec1 were determined to be fragments and removed. This threshold was selected based on manual inspection of the aligned structures. The resulting dataset included 23,082 total sequences, including both eukaryotic and prokaryotic CLC sequences.

Additional curation of sequences was performed for the various alignment strategies used in the phylogenetic analysis described below.

### Phylogenetic analysis and CLC functional classification

Different sequence alignment strategies were tested to examine the robustness of the phylogenetic analysis. For each method, an approximate maximum likelihood tree was calculated with FastTree using the LG model, 100 traditional bootstraps and other default parameters. Support values were calculated along key nodes in the CLC phylogeny, and the sum of support values was calculated over all the nodes on a dataset consisting of all eukaryotic homologs, including those found outside the main clade. All alignment methods used a pairwise structural alignment to a defined reference structure. We compared alignments generated from altering the following variables (**Supp. Table 2**): (1) reference structures of AF2 models of either CLC-ec1 (P37019) or hCLC-2 (P51788), (2) alignment site definitions corresponding to residues in transmembrane alpha helices or all residues with alignment occupancies > than 85%, (3) sequence alignment correction with MUSCLE v3, and (4) including only the transmembrane core or including intracellular domains CBS1 and CBS2. The key nodes examined were: node 1 - the most recent common ancestor (MRCA) of hCLC-1 and hCLC-K (62 on the final tree), node 2 - the inner flanking node of the TG (96 on the final tree), node 3 - the outer flanking node of the TG (96 on the final tree), and node 4 - the MRCA of the outgroup sequences (N/A on the final tree, only 1 outgroup sequence used). We found that hCLC-2 was a better reference structure than CLC-ec1, most likely because the alignment to eukaryotic homologs has higher overall occupancy, providing more information to build a tree. Using residues outside of the CLC TM core showed no clear improvement. In fact, adding the rest of the sequence to the MSA removes node 3 entirely despite having a high support value in all other trees. Among the alignment strategies that used only the TM core, including the high occupancy alignments sites outperformed using only TM residues. Lastly, sequence alignment correction had little effect on the total support values, but improved nodes 1-3. Therefore, the alignment used in this study included all alignment site occupancies > 85% in the TM core, hCLC-2 as the reference structure and a sequence alignment correction.

An initial phylogenetic analysis was done on the set of sequences clustered at 80% sequence identity in the TM regions, retaining one representative per cluster. In tota, there were 10,417 sequences, including both eukaryotic and prokaryotic homologs. The tree was generated using FastTree (LG, gamma only for final branch lengths, 100 bootstraps) using an MSA generated from a near full-length structure alignment with CLC-ec1. For the second phylogenetic analysis, the tree was generated using RAxML (LG+G, 100 bootstraps), and the input MSA was generated from a near full-length structure alignment with hCLC-2. The sequence alignment tool MUSCLE was then used to correct for possible frameshifting in the structure alignment; we used the raw structure alignment as input with default settings. Phylogenetic trees were visualized on iToL (Letunic & Bork, 2021). To improve the tree visuals, the outgroup branch length was manually set to 0.1 in Figure 4.

To classify sequences by putative function, we make one of two assumptions. First, we define an arbitrary evolutionary distance cutoff indicating that the functional switch did not occur in recent evolutionary history. For this, a sequence was classified as belonging to a certain function if that sequence was less than 1.5 evolutionary units away from a reference sequence of that known function in both the final tree and at least 85% of bootstrap support trees. Second, we assume that the functional switch occurred only once. To define the transition group, we note that sequences cmCLC (transporter (Feng et al., 2010a)) and ngCLC (channel (Fortea Verdejo, 2021)) were found in a low-resolution region of the tree, situated between two nodes with 96% support values (see **Fig. 4F**). Because this region contains both channel and transporter homologs in proximity, we conclude that the functional switch occurred here (as was concluded previously (Fortea Verdejo, 2021)). Based on the assumption that only one functional switch exists, all sequences inside of the inner flanking node are classified as channels and all sequences outside of the outer flanking node are noted as transporters. Due to their proximity to the functional switch, none of the sequences within the transition group itself were classified as either transporter or channel and instead treated as a separate CLC subfamily defined as the transition group (TG).

### Distance and Distance Difference Matrix Analysis

To compare different CLC structures, we carried out pairwise Cα atom structural alignments of the 1266 eukaryotic AF2 CLC models to the reference CLC-ec1 (AF2 model, Uniprot ID: P37019) using TM-align. Positions within the transmembrane helices of CLC-ec1 that exhibited 85% occupancy across this CLC set were selected to build a common list of alignment sites. Residues in the αC-D linker (100-109) and helix R (444-450) that are known to contain key functional residues (e.g., S107, Y445) were added for a total list of 1-294 alignment sites (**Supp. Table 3**).

A 3D protein structure consists of a set of atom positions in Cartesian coordinates, with each residue containing a Cα backbone atom. A CLC structure is numerically represented as a distance matrix, DM(*i*, *j*), containing the scalar distances between Cα atoms for all pairs of residues (*i, j*) within the structure:

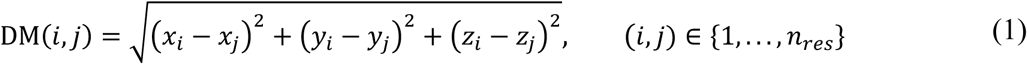

To compare between CLC homologs, we define a common subset of residues defined as alignment sites and trim the distance matrix to yield the aligned distance matrix DM_aln_(*m, n*):

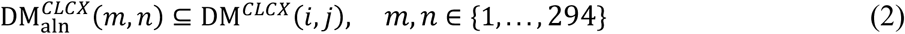

where (*i*, *j*) are the residue indices corresponding to alignment sites (*m*, *n*) on CLC-X. For a given CLC sequence/structure, it is possible that a residue does not exist for the corresponding alignment position (i.e., gaps in the structure alignment); in this case, DM_aln_(*m,n*) is set to the null value of “*NaN*”, i.e., “*Not a Number*”.

Distance difference matrices (DDM) represent the absolute distance differences between two structures or groups of structures. To quantify the structural differences between any two CLC sequences, e.g., CLC-X and CLC-Y, we calculate:

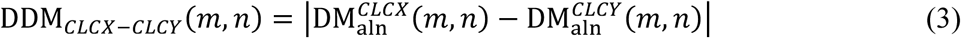

Further, DDMs between two groups of structures can be calculated, such as between all structures associated with one gene type, e.g., human CLC-1 and human CLC-7 or all CLCs classified as transporters vs. channels. The DDM between groups of structures A and B, represent the difference between the averages of all CLCs in that group:

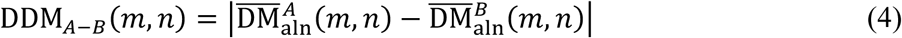

Here, 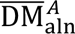 represents the average distance matrix for all structures in a group A:

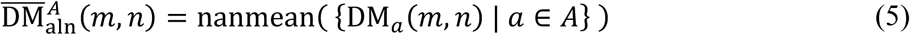

and likewise for group B. Note that ‘nan’ operations like nanmean ignore all *NaN* values in the matrix. Heatmaps of DMs and DDMs are made using the Python visualization library Matplotlib (Hunter, 2007). Helix breaks are designated as rows or columns of *NaN* values, ensuring no data is obscured.

The mean across all distance differences associated with each residue *m* yields a mean distance difference line plot, i.e., mDD_res._(*m*).

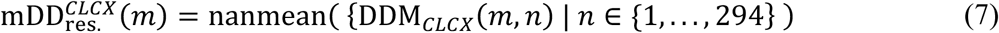

If there is no residue at *m* because of a gap in the structure alignment, then mDD_res._ is set to *NaN*. For line plots comparing between two groups of structures, e.g., all channel and transporter structures, we compare between the averages of all *mDD_res._(m)* line plots in each group (**Supp. Fig. 2C**):

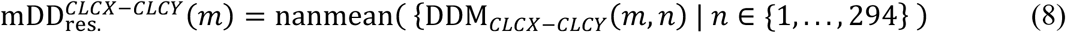

A global average of distance differences over the whole structure, mDD_struct._ provides a quantitative metric to summarize how different two structures are, like an RMSD for a pairwise structure alignment, by averaging over the entire DDM:

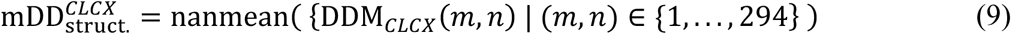

### Ensemble-based model-fitting

To identify structural features that distinguish CLC transporters from channels, we employed a two-step model-fitting approach. First, we trained a Random Forest Classifier (RFC) to discriminate between structural representations of predicted channel and transporter models, using flattened distance matrices, i.e., a vector of length *l* = *m x n*, containing all pairwise distances as input features (**Fig. 4A**). We selected the RFC due to its ability to capture nonlinear relationships while allowing for efficient computation of SHAP values, a critical requirement for interpreting model decisions. In principle, other classifiers could be substituted to trade off interpretive power or computational cost, but RFCs offered a favorable balance for our large dataset. RFCs were trained using the Python machine-learning library *Scikit-Learn* (Pedregosa et al., 2018) on selected transporter and channel structures with the following parameters: n_estimators = 500, max_depth = 5, max_samples = 0.3. The output of the RFC is the probability that a particular CLC sequence is a channel, *P_Chan_*, determined by the classifier’s learning of structural features associated with channel or transporter groups.

Second, we analyzed the trained model to quantify the average expected marginal contribution of an input feature to the calculated channel probability using SHAP values. SHAP values represent the “importance” of an input feature to a classifier’s prediction. It is denoted as *SHAP(m,n,s,r)*, where *(m,n)* = {(1, …, 261), (1, …, 261)} alignment site input features, *s* = {1, …, 1620} structures in this analysis, and *r* is the RFC iteration. For input features that do not exist, i.e., gaps in a structure, *SHAP(m,n,s,r)* is manually set to NaN prior to the training of that RFC. All other *SHAP(m,n,s,r)* calculations are carried out using the SHAP command in *Scikit-Learn* (S. Lundberg & Lee, 2017).

Let *T* be the set of all structures *s* in the training set. Then, the absolute SHAP value, i.e., *absSHAP*, is calculated as an average over all structures to determine the importance of a single feature site to an RFC:

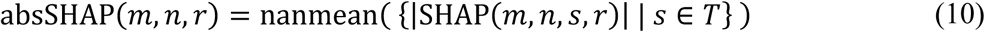

The RFC analysis procedure required further optimization to identify the robust structural changes between the two groups of CLCs. Initial trials with a single RFC revealed overfitting to a sparse subset of features. For instance, once the model learned that a specific pairwise distance was predictive, e.g., *d_m,n_*, it tended to assign near-zero importance to neighboring correlated features, i.e., *d_m+1,n_*. This led to poor feature sampling and skewed SHAP distributions dominated by a few outliers. To mitigate this, we tested two ensemble approaches where we iterate the RFC classification to analyze over the entire RFC population and also examine a masking procedure where each RFC is trained on input data with a randomly masked subset of features. We define the overall importance value as the average of the absSHAP values over *r* separate RFCs. Let *R* be the set of all trained RFCs:

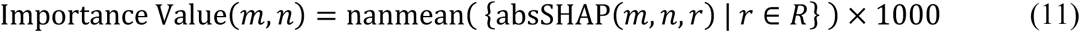

where 1000 is added as an arbitrary scaling factor to simplify plotting. Increasing the number of RFC replicates addressing the overfitting by producing ‘smoother’ importance value matrices (**Supp. Fig. 6A**). The overall distribution of importance values also spreads out, making it clearer to separate out the structural features associated with the strongest importance values (**Supp. Fig. 6B**). Based on these results, RFCs were analyzed over 1000 iterations in this study.

In the case of masking, a fixed percentage of features were excluded from the input data prior to training of an RFC. This stochastic masking procedure forces individual models to rely on different subsets of the input space, improving the uniformity of SHAP value distribution and enabling the detection of weaker but biologically relevant signals. However, we found that there was not a strong dependency of the importance values on the percentage of masked features as they generally yielded similar distributions up to 60% masking (**Supp. Fig. 6C, D**). In addition, masking does not significantly change the output probabilities of the RFCs relative to unmasked until 60% (**Supp. Fig. 6E**). This is likely because RFCs internally introduce differential sampling of data for each branch of its tree, so the masking is redundant. Loss of information in the importance value distribution was observed at 80% masking. Based on these results, we conclude that masking is not a required step in this classification procedure but suggest that it be tested if other deterministic classifiers are used in place of RFCs. In the end, a masking level of 20% was adopted for our analysis in this paper.

### Structural analysis of CLCs

A list of experimentally determined CLC structures containing the transmembrane domains were identified from the RCSB Protein Databank (**Supp. Table 1**) and DDMs relative to CLC-ec1 (PDB ID: 1OTS) were calculated as described previously (**Supp. Fig. 1**). AF2 structures were downloaded from the AlphaFold Protein Structure Database (Varadi et al., 2024) following the list of Uniprot IDs for the IPR001807 family of CLC proteins. Histograms of the absolute distance differences (DD) were calculated between -0.01 to 5.99 Å with bin width 0.2 Å and fit with a Gaussian function in GraphPad Prism v10.4.0:

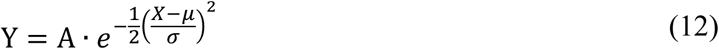

where *A* is the amplitude, *μ* is the mean and *σ* is the standard deviation. When clustering structural features on a DDM, groupings were selected according to unique helix pairs and then combined into larger clusters hypothesized to describe a common structural change.

The 3D structures of hCLC-1 (PDB ID: 6COY) and hCLC-7 (PDB ID: 8HVT) were compared as representative experimental examples showing maximal changes in regions of high importance. For visualization, structures were aligned using TM-align and then the region of interest was highlighted onto the transmembrane domain of the hCLC-1 structure. To create a ‘morph’ movie showing the linear interpolation between the AF2 models (**Supp. Video 1**), a pair of AF2 models was chosen to highlight the changes with highest importance and minimize all other extraneous changes. We screened all possible eukaryotic structure pairs, calculating the average distance difference for distances in the top 5% and bottom 90% of importance values. From this, we selected bsCLC from *Betta splendens* as the channel candidate (Uniprot ID: A0A6P7PB93) and cfCLC from *Cavenderia fasciculata* as the transporter candidate (Uniprot ID: F4PTB4), whose average distance difference for the top 5% in importance is 1.57 Å and average distance difference for the bottom 10% is 0.49 Å (**Supp. Fig. 5**). The transporter structure was structurally aligned to the channel structure with TM-align; residues that did not structurally align were removed, while residues that aligned were mutated on the transporter structure to be equivalent to the channel structure, creating two structures with one-to-one atomic correspondence. A linear morph between the two models was created and visualized using VMD (Humphrey et al., 1996).

## SUPPLEMENTAL INFORMATION

### Distance difference matrices show variation amongst experimental CLC structures

Existing experimental structures show structural variation relative to CLC-ec1 (**Supp. Fig. 1**). As mutations accumulate, sequence variation eventually results in changes in a structure’s pairwise distances. For example, stCLC shares ∼80% sequence identity with CLC-ec1 and shows a DDM with modest structural changes with 73% of distance differences < 0.5 Å and only 1% > 3 Å (mDD_struct._ = 0.50 Å). CLC-sy1 only shares ∼40% sequence identity but is still quite similar in structure to CLC-ec1 (58% DDM values < 0.5 Å, 1% > 3 Å; mDD_struct._ = 0.68 Å). On the other hand, while CLC-F is prokaryotic, it is the most different from CLC-ec1 (mDD_struct._ = 1.39 Å) with 32% of DDM values < 0.5 Å and 11% > 3 Å, exceeding the changes observed in the eukaryotic CLC structures. In general, the structural differences observed correspond to the evolutionary distance of these sequences relative to CLC-ec1 (**Fig. 4B**) and demonstrate the structural variability obtained in the different experimental CLC structures. We note that while CLC-F and the eukaryotic CLCs are similarly different to CLC-ec1, they are also similarly different to each other (e.g., mDD_struct._ = 1.60 Å between CLC-F & hCLC-2). Importantly, this analysis demarcates the limits of how different CLC structures can be while maintaining the same topological architecture and fold.

### Validating CLC models predicted by AlphaFold2

AF2 was trained on unique structures in the PDB before April 30, 2018, meaning that about half of the experimental CLC structures available today were not in the original training set. To investigate whether AF2 can successfully predict CLCs that were not in the training set, and report on structural differences between different CLCs, we calculated DDMs for each experimentally determined eukaryotic CLC and the AF2 prediction for the same CLC gene. For most eukaryotic CLCs, there are multiple experimental structures reflecting different conformations, so we examined each PDB separately and plotted the DDMs for the most and least similar experimental structures (**Supp. Fig. 2A**). Comparing the different PDBs, we see that there are substantial differences between experimental and predicted structures; however, there is typically one PDB that agrees well with the AF2 prediction such that their mDD_struct._ < 0.5 Å, on the order of what we observed when comparing similar sequences such as stCLC and CLC-ec1 (**Supp. Fig. 1**). For example, both structures of hCLC-2 (PDB ID: 7XJA & 8TA3) are like their AF2 prediction with mDD_struct._ = 0.23 Å and 0.44 Å, respectively. With this, we conclude that AF2 predicts the structure of different CLC genes accurately overall and within the range of distance differences that would be expected for two similar sequences.

To examine whether AF2 successfully predicts all regions of CLC structure, we examine the mDD_res._ line plots (**Supp. Fig. 2B**). Again, we see that there is always one PDB structure that reflects an accurate prediction, but this analysis also highlights regions that are not well predicted by AF2. Averaging over the 15 mean absolute distance difference line plots gives us a global mean and standard deviation, revealing that the structural elements that are less accurately predicted are the ends of helix αH and αF (**Supp. Fig. 2C**). These deviations arise from hCLC-1 as well as the transporter homologs hCLC-6 & hCLC-7, where AF2 underpredicts the distance between helix αH and the rest of the structure, placing helix αH closer to the central core of the subunit than what is observed in the experimental structures. Similarly, for hCLC-6, hCLC-7, and plant transporter atCLCa, AF2 tends to position the end of helix αF closer to the protein core. Other deviations also exist but appear to be unique to a certain CLC type. For instance, examining the three structures of hCLC-1 channel, we see that PDB IDs 6QVD and 6QV6 show the largest deviations in the αC-D linker, where S189 along the transport pathway is flipped in one structure, and is not predicted by AF2. However, this serine flipping is not observed in all hCLC-1 structures (e.g. PDB ID 6COY), indicating it is likely just a conformational state. hCLC-2 also shows a unique deviation in helix αI that is not present in the other CLCs. These unique deviations tend to disappear in the averaging of the structural changes leaving the ends of helix αH and αF as the most prevalent deviations in the predicted structures observed in about half of the structures analyzed. At large scales, however, we hypothesize that these changes average out and become statistical noise in the analysis.

Another way of assessing the quality of an AF2 prediction is the predicted local distance difference test (pLDDT) score, which represents AlphaFold’s confidence in the placement of each residue relative to its surrounding residues. The pLDDT scores are plotted for each residue position alongside the mean absolute distance difference per residue (**Supp. Fig. 2B**). Deflections are observed in the pLDDT scores, but ∼80% of residues analyzed exhibit pLDDT values > 90 (**Supp. Fig. 2D**), indicating high confidence in all CLC predictions. We note that the pLDDT scores do not show correlations between mean distance differences (**Supp. Fig. 2D**), so regions where we observe structural deviations (e.g., the αC-D linker, helix αH and αF) do not always correspond to low pLDDT values. We plotted the mDD_struct._ for all experimental-AF2 DDMs and existing experimental structures to CLC-ec1 (**Supp. Fig. 2E**). Among these, prokaryotic transporters CLC-F and CLC-ec1 are the best predicted structures by AF2. The eukaryotic channel hCLC-1 is least successfully predicted, but this is also the construct that exhibits the largest conformational variability among experimental structures, with mDD_struct._ scores ranging from 0.4-1.1 Å. In general, AF2 predictions for sequences included in the training set are comparable to sequences from outside the training set, and all AF2 predictions yield smaller mDD_struct._ scores obtained when comparing the experimental differences between two CLC genes, even those that are similar in sequence like CLC-ec1 and stCLC. These results support that AF2 successfully predicts a correct structure corresponding to one accessible conformation for each CLC gene.

While the accuracy of AF2 predictions may deviate in certain structural regions, it is possible that these errors are systematic. In this case, even if individual models deviate from the experiment data, AF2 may still be able to preserve *relative* structural differences between CLC homologs. In other words, AF2 could be more reliable at capturing structural variation across sequences than in recapitulating any one structure precisely. Since we are ultimately interested in identifying changes between different groups of CLCs, we evaluated whether AF2 can reliably capture relative structural differences between homologs. To inspect this, we compared pairs of CLC homologs, calculating DDMs between their experimental structures and then between the respective AF2 models (**Supp. Fig. 3A**). Examining the mDD_res._ line plots, we can see that there is a clear correlation between the predicted differences from AF2 and those that are experimentally observed (**Supp. Fig. 3B**). Notably, in the hCLC-1 vs. hCLC-7 comparison, structural differences in helix αH are accurately reflected in the AF2 DDM even though AF2 misplaces the end of helix αH in both individual models (**Supp. Fig. 2B**). The most significant underprediction in this comparison occurs at the αC-D linker, due to the flipped serine conformation present in the 6QV6 structure that is not well captured by AF2. In the hCLC-2 vs. hCLC-6 comparison, helix αH exhibits some of the largest predicted differences between the two structures, yet the magnitude remains underestimated relative to experimental data. On the other hand, homolog pairs with the same function show more subtle structural differences overall. In the hCLC-6 vs. hCLC-7 comparison, AF2 captures the key structural change at helix αF between these transporters, albeit with reduced amplitude. Even more subtle differences, such as in helix αC, are weakly but detectably reflected in the predictions. In stark contrast, AF2 predicts virtually no structural differences in hCLC-1 vs. hCLC-2 (mDD_struct._ = 0.21 Å), despite the experimental DDM indicating significant changes (mDD_struct._ = 0.59 Å). The reason for this discrepancy remains unclear; the two sequences share a moderate sequence identity of ∼70% across the DM_aln_ alignment sites, like the identity between bCLC-K and hCLC-2 (∼53%) which does not exhibit the same level of underprediction. To quantify the correlation and underprediction, we calculate DDMs for all possible homolog pairs among those with known structures and plot mDD_res._ calculated from experiment and AF2 against each other (**Supp. Fig. 3C**). The line of best fit has a slope of 0.56 and R^2^ = 0.60 indicating that AF2 predicts distance differences but is reduced by a factor of 1.8.

Finally, we compare the calculation of a DDM between channels and transporters using only the experimental structures available and the corresponding AF2 models so that we examine the same group of structures (**Supp. Fig. 4A-C**). Following the SHAP importance analysis, we isolate the largest values from these DDMs, in this case representing the largest structural differences by magnitude of the change, plotting the top 1% and 2% of the distance differences in each DDM (**Supp. Fig. 4D-F**). Consistent with the DDM using experimental structures, AF2 predicted that the largest structural differences were concentrated in helices αF, αH, αJ, and αK (**Supp. Fig. 4G-I**). A notable divergence from the experimental results was the absence of large distance differences involving the αC-D linker (**Supp. Fig. 4J**). Since the flipping of the αC-D linker is a dynamic feature associated with the active or open state of the protein, it reflects a conformational change rather than a more permanent structural feature associated with the protein’s fold. In place of the αC-D linker, the AF2 DDM reveals large distance differences between helix αH and the N-terminal half of the protein, as well as within helix αP. These changes appear in the top 1% of the experimental DDM values and replace the distance differences vacated by the αC-D linker in the AF2 DDM.

From this, we conclude that AF2 can predict both the overall structure and structural differences of CLC homologs. While individual predictions may contain inaccuracies, differences are aggregated across thousands of structures, significantly improving the signal to noise ratio. Our results do, however, reveal a systematic underestimation in distance difference magnitudes predicted from AF2 by a factor of 1.8. We find that while AF2 preserves observed changes, the extent of those changes are conservative, likely due to a bias toward predicting the most stable, “average” conformation to minimize prediction error. AF2-derived distance differences should therefore be interpreted as lower bounds for the true extent of change.

## SUPPLEMENTARY FIGURES

**Supplementary Figure 1.**
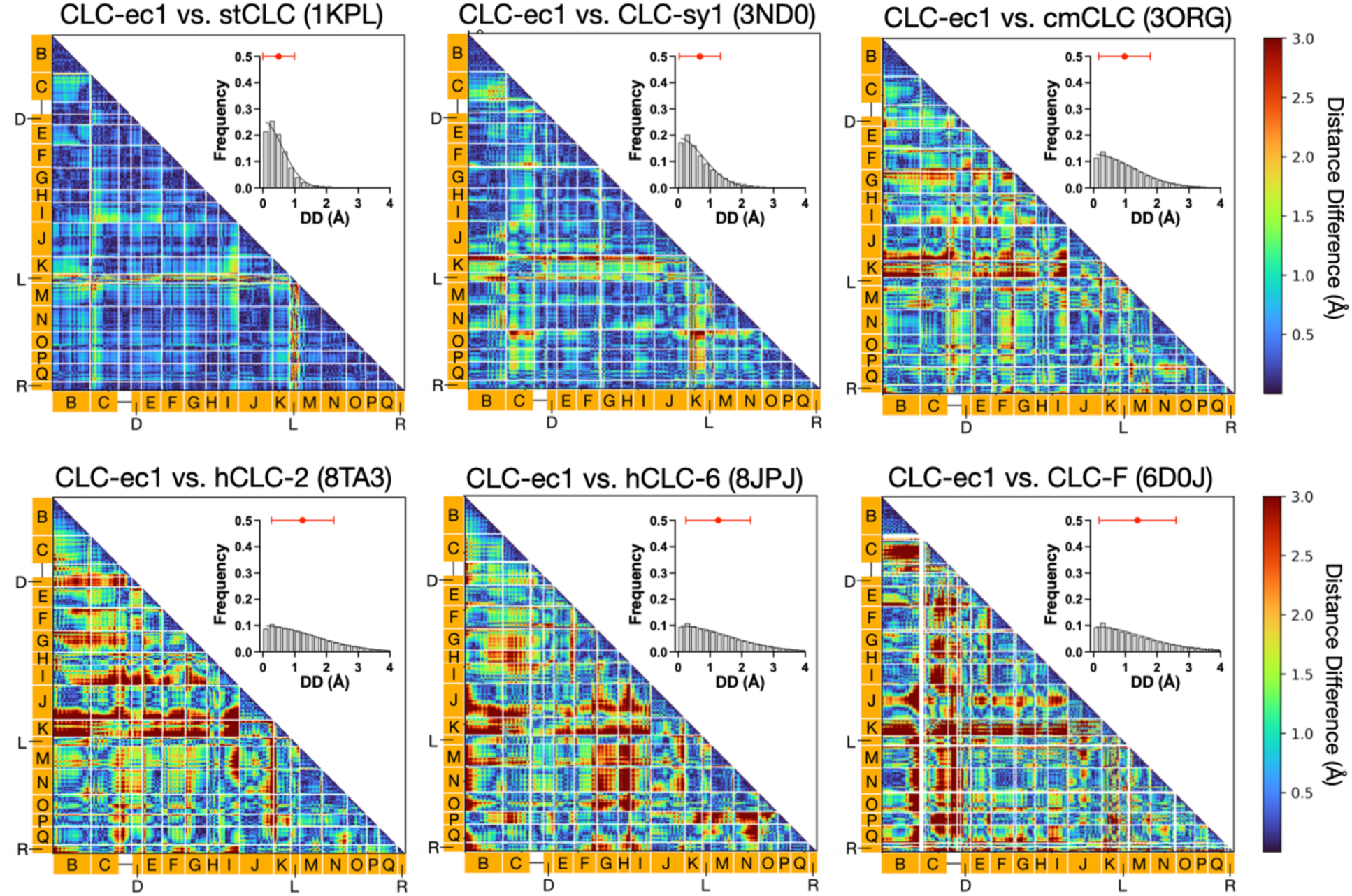
Individual DDMs of experimental CLC homolog structures compared to CLC-ec1. Absolute distance difference matrix (DDM) maps for experimental structures of different CLC homologs compared to the CLC-ec1 (PDB ID: 1OTS) reference structure. Frequency histograms of the absolute distance differences are shown in the inset plots, with distributions fit by Gaussian functions centered at μ = 0 (black line). The mean ± std of all distance differences are also shown (red point). Plots are ordered according to greatest to least similarity to CLC-ec1: prokaryotic stCLC (PDB ID: 1KPL, A = 0.30, σ = 0.52, mean ± std = 0.50 ± 0.50) and CLC-sy1 (PDB ID: 3ND0, A = 0.21, σ = 0.71, mean ± std = 0.68 ± 0.65), eukaryotic cmCLC (PDB ID: 3ORG, A = 0.14, σ = 1.14, mean ± std = 0.98 ± 0.81), hCLC-2 (PDB ID: 8TA3, A = 0.10, σ = 1.51, mean ± std = 1.25 ± 1.00), and hCLC-6 (PDB ID: 8JPJ, A = 0.10, σ = 1.52, mean ± std = 1.32 ± 1.26), and prokaryotic CLC-F (PDB ID: 6D0J, A = 0.10, σ = 1.50, mean ± std = 1.44 ± 1.32) .

**Supplementary Figure 2.**
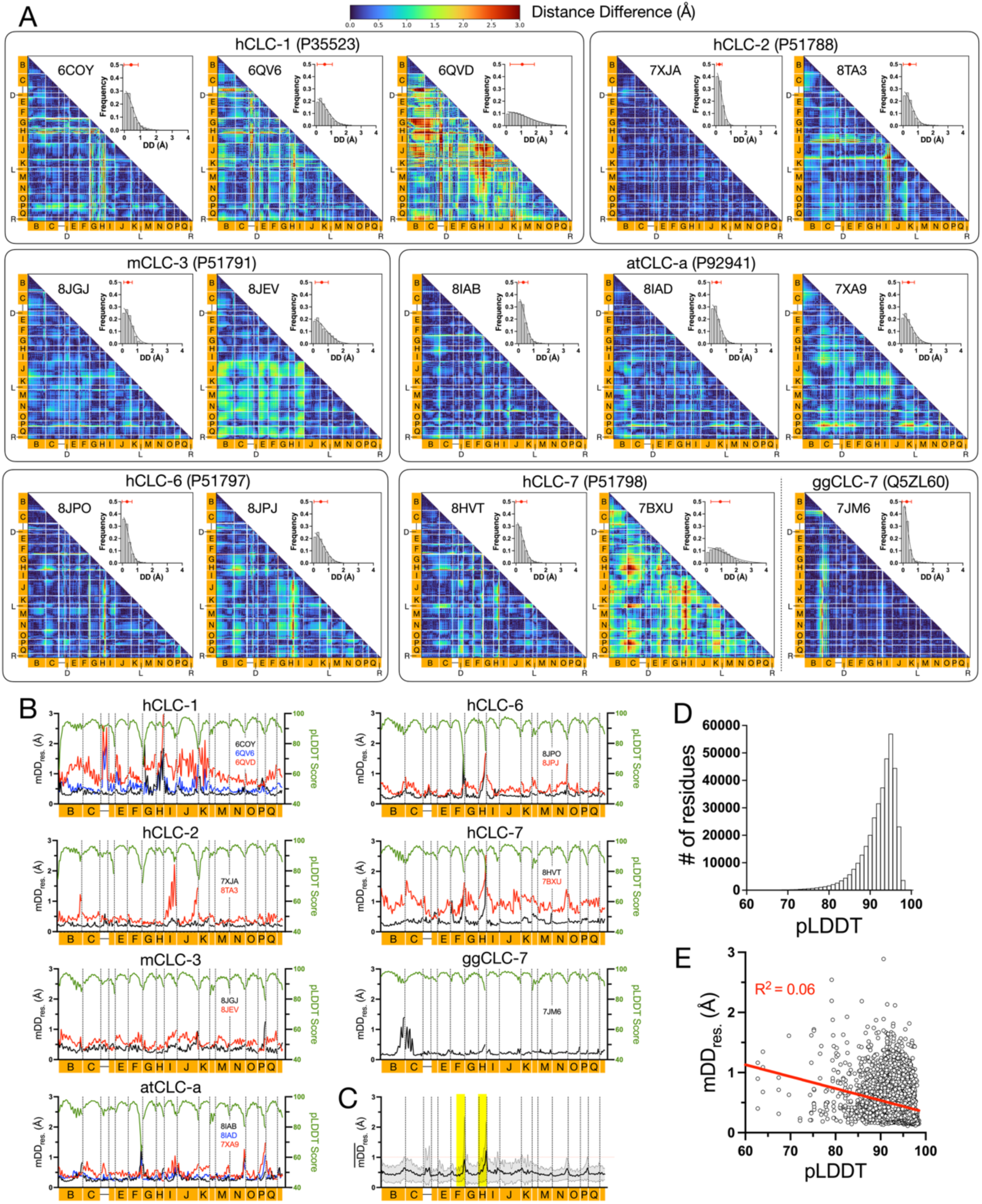
AlphaFold2 vs. experimental structure distance difference matrices. **(A)** Absolute DDMs for experimentally solved CLC structures since 2018 compared to its AF2 model. For each gene, when multiple structures have been solved, absolute DDMs are ranked left to right from most to least similarity. **(B)** Mean absolute distance differences for each residue for each PDB (black, blue & red) for each CLC gene, alongside pLDDT scores (green). **(C)** mDD_res._ ± standard deviation over 15 structures examined in (B). Yellow regions indicate structural elements with deviations > 1 Å. **(D)** Histogram of pLDDT values over all residues. **(E)** Mean absolute distance differences per residue vs. pLDDT. Linear regression line (red) shows no correlation (R^2^ = 0.06). **(F)** Mean absolute distance differences per structure for AF2 models vs. PDBs in the training set (red), AF2 models vs. PDBs determined since 2018 (blue), and experimental PDBs vs. CLC-ec1 PDB ID: 1OTS from **Supp.** Fig. 1 (grey).

**Supplemental Figure 3.**
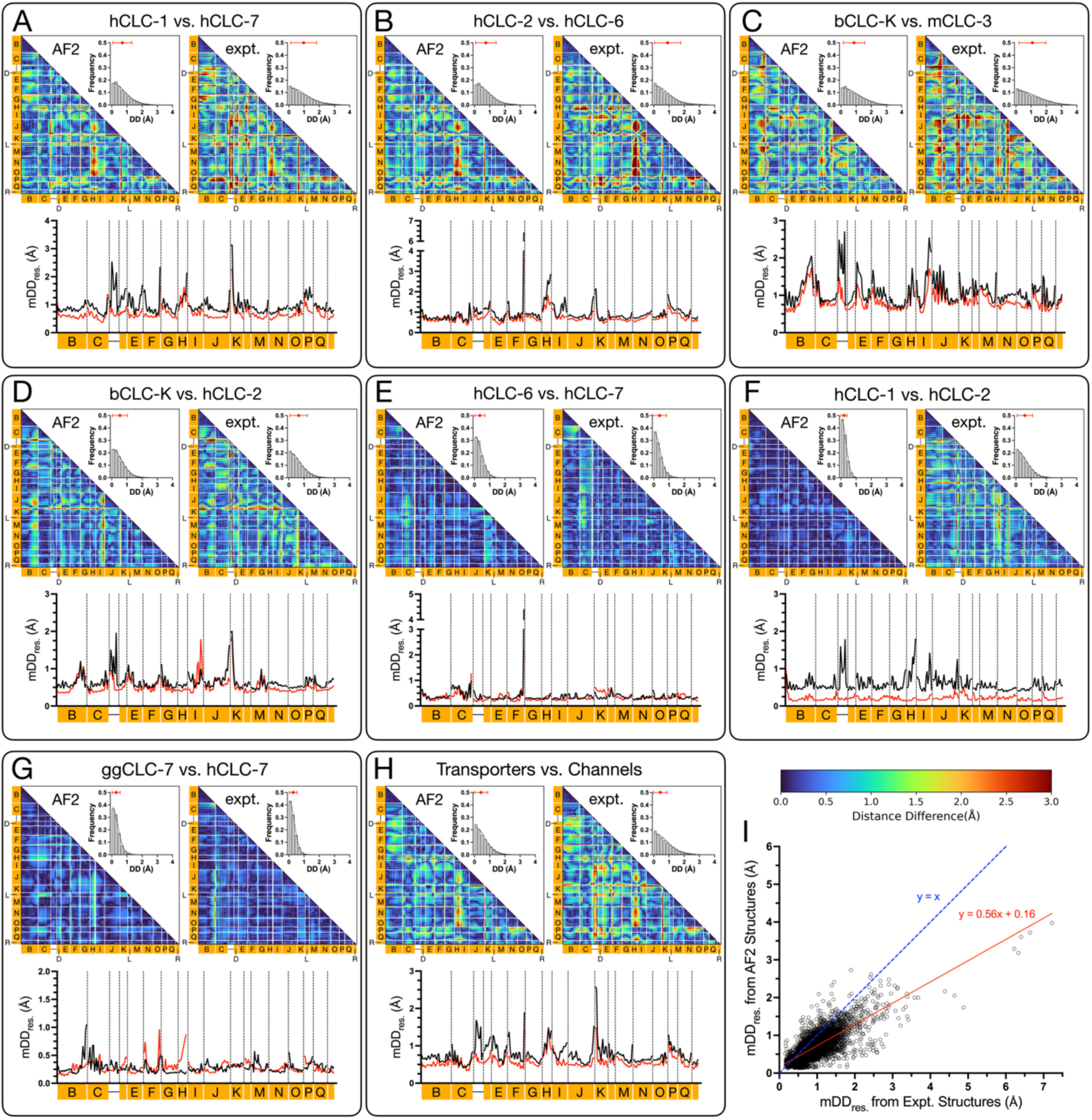
Relative differences between AF2 and experimental structures. **(A-H)** For each pair of CLC homolog groups the DDMs, DD histograms, mDD_struct._ and mDD_res._ line plots (experimental structures in black, AF2 models in red) are shown. (**I)** Scatterplot between the mean distance differences by residue for AF2 predictions compared to experimental structures. Linear regression analysis shows the best-fit line (R^2^ = 0.60) with slope of 0.56 and y-intercept = 0.16 (red).

**Supplementary Figure 4.**
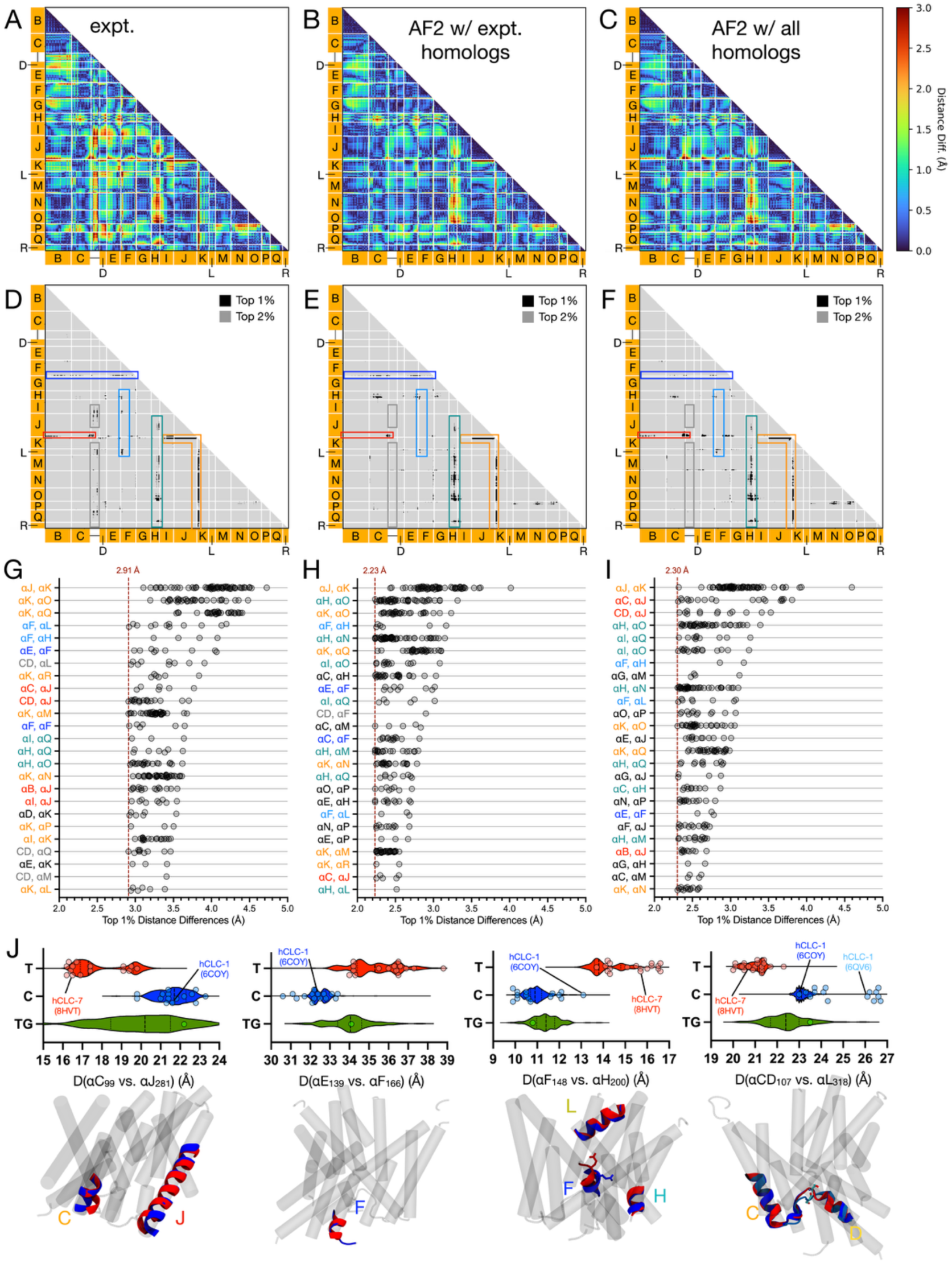
Channel vs. Transporter DDMs with different structure selections. **(A-C)** DDMs between channels and transporters using experimental structures, corresponding AF2 models for the experimental structures, and all available AF2 models, respectively. **(D-F)** Following the SHAP importance analysis, the top 1% and 2% of distance differences are shown to demonstrate the largest structural changes for each set of structures used. **(G-I)** Top 1% distance differences plotted based on their location on the corresponding DDM; each segment of the DDM is ranked based on its maximum distance difference. **(J)** Structural depictions and histogram plots for the distances between residue pairs that are large but do not show up as highly important from the SHAP analysis (**Fig. 5**).

**Supplementary Figure 5.**
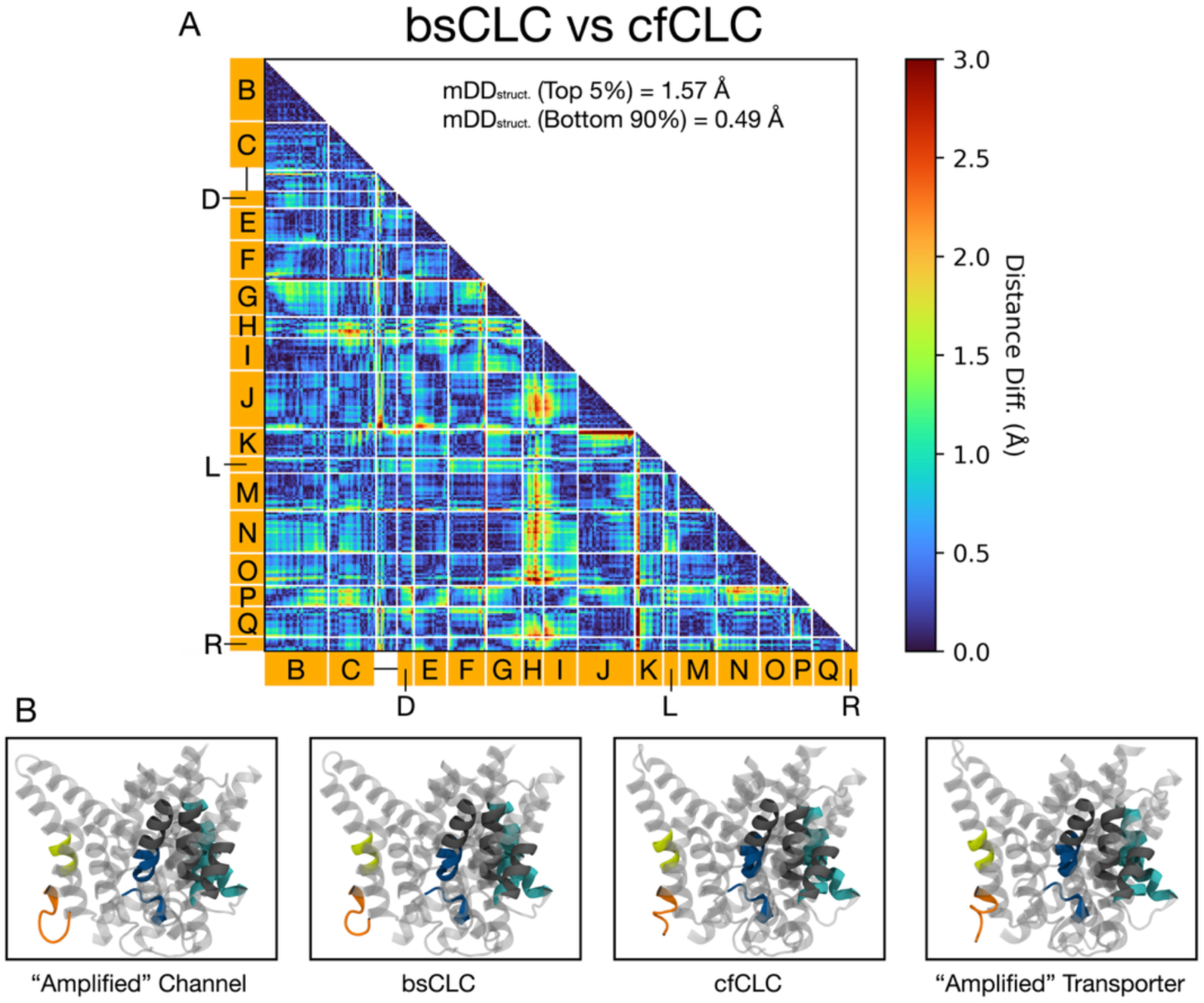
A comparison of CLC channel and transporter structural changes. **(A)** DDM between the channel bsCLC (A0A6P7PB93) and transporter cfCLC (F4PTB4), as a representative example of the structural changes observed. These homologues were selected from the overall mDDM between all AF2 transporters and channels, taking the top 2% importance values, and bottom 50% importance values, and selecting a transporter/channel DDM by eye that captures the high importance value features and not the lowest. **(B)** Example frames from a morphing video between bsCLC and cfCLC models. From left to right, amplified channel, channel, transporter, amplified transporter.

**Supplementary Figure 6.**
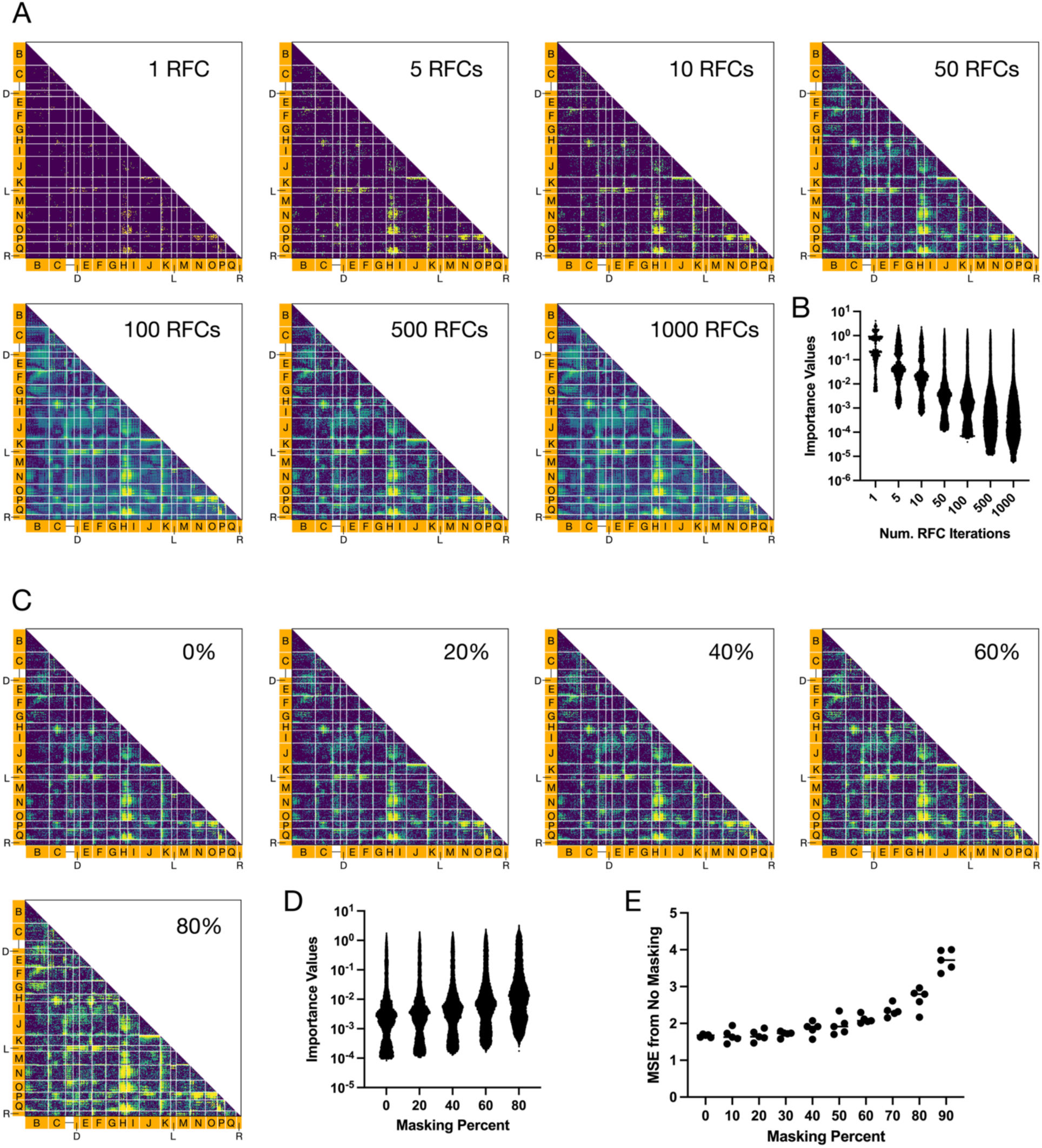
Optimization of the ensemble importance value calculation. **(A)** Average importance value matrices as a function of random forest classifier (RFC) replicates with 20% masking. **(B)** Importance value distributions as a function of RFC replicates. **(C)** Average importance value matrices for 50 RFC replicates as a function of masking percentage. **(D)** Importance value distributions for the corresponding plots, demonstrating little to no change based on masking percent. **(E)** Effect of masking percent on output probabilities of the RFCs (5 replicates each). On the y-axis is the root mean squared error from the probability outputs of a single RFC with no masking.

**Supplementary Video 1. Linear interpolation between channel bsCLC and transporter cfCLC**. A morph video depicting the structural transition between channel bsCLC (A0A6P7PB93) and transporter cfCLC (F4PTB4) by linear interpolation. Structures were aligned with TM-align, creating one-to-one atomic correspondence by mutating cfCLC to be identical to bsCLC with pymol mutagenesis, and a modified morph script in VMD generating frames past the true transporter and channel structures to amplify the structural differences.

## SUPPLEMENTARY TABLES

**Supplementary Table 1:**
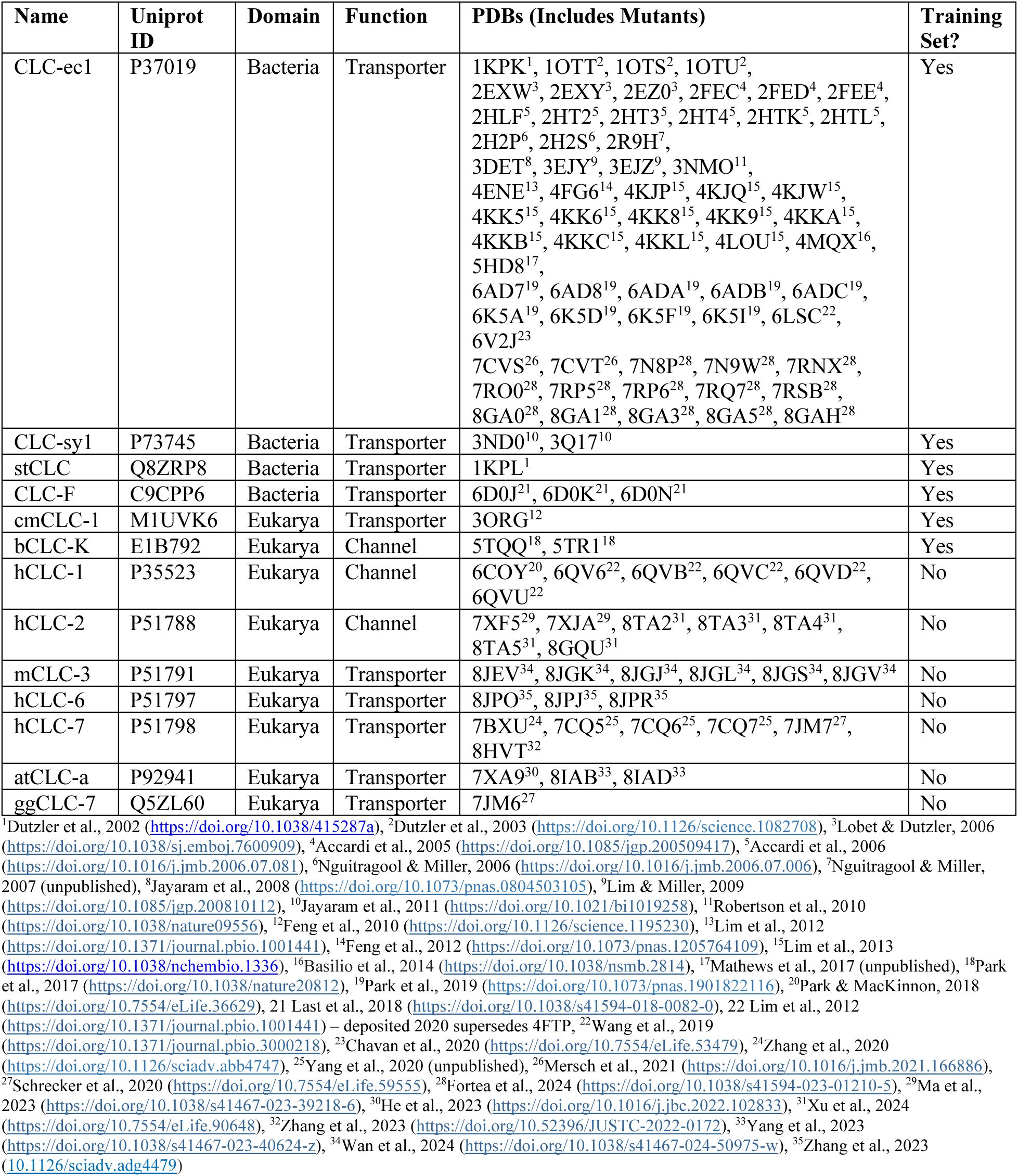
List of all experimental CLC Structures in the PDB.

| Name | Uniprot ID | Domain | Function | PDBs (Includes Mutants) | Training Set? |
| --- | --- | --- | --- | --- | --- |
| CLC-ec1 | P37019 | Bacteria | Transporter | 1KPK<sup>1</sup>, 1OTT<sup>2</sup>, 1OTS<sup>2</sup>, 1OTU<sup>2</sup>, 2EXW<sup>3</sup>, 2EXY<sup>3</sup>, 2EZ0<sup>3</sup>, 2FEC<sup>4</sup>, 2FED<sup>4</sup>, 2FEE<sup>4</sup>, 2HLF<sup>5</sup>, 2HT2<sup>5</sup>, 2HT3<sup>5</sup>, 2HT4<sup>5</sup>, 2HTK<sup>5</sup>, 2HTL<sup>5</sup>, 2H2P<sup>6</sup>, 2H2S<sup>6</sup>, 2R9H<sup>7</sup>, 3DET<sup>8</sup>, 3EJY<sup>9</sup>, 3EJZ<sup>9</sup>, 3NMO<sup>11</sup>, 4ENE<sup>13</sup>, 4FG6<sup>14</sup>, 4KJP<sup>15</sup>, 4KJQ<sup>15</sup>, 4KJW<sup>15</sup>, 4KK5<sup>15</sup>, 4KK6<sup>15</sup>, 4KK8<sup>15</sup>, 4KK9<sup>15</sup>, 4KKA<sup>15</sup>, 4KKB<sup>15</sup>, 4KKC<sup>15</sup>, 4KKL<sup>15</sup>, 4LOU<sup>15</sup>, 4MQX<sup>16</sup>, 5HD8<sup>17</sup>, 6AD7<sup>19</sup>, 6AD8<sup>19</sup>, 6ADA<sup>19</sup>, 6ADB<sup>19</sup>, 6ADC<sup>19</sup>, 6K5A<sup>19</sup>, 6K5D<sup>19</sup>, 6K5F<sup>19</sup>, 6K5I<sup>19</sup>, 6LSC<sup>22</sup>, 6V2J<sup>23</sup>, 7CVS<sup>26</sup>, 7CVT<sup>26</sup>, 7N8P<sup>28</sup>, 7N9W<sup>28</sup>, 7RNX<sup>28</sup>, 7RO0<sup>28</sup>, 7RP5<sup>28</sup>, 7RP6<sup>28</sup>, 7RQ7<sup>28</sup>, 7RSB<sup>28</sup>, 8GA0<sup>28</sup>, 8GA1<sup>28</sup>, 8GA3<sup>28</sup>, 8GA5<sup>28</sup>, 8GAH<sup>28</sup> | Yes |
| CLC-sy1 | P73745 | Bacteria | Transporter | 3ND0<sup>10</sup>, 3Q17<sup>10</sup> | Yes |
| stCLC | Q8ZRP8 | Bacteria | Transporter | 1KPL<sup>1</sup> | Yes |
| CLC-F | C9CPP6 | Bacteria | Transporter | 6D0J<sup>21</sup>, 6D0K<sup>21</sup>, 6D0N<sup>21</sup> | Yes |
| cmCLC-1 | M1UVK6 | Eukarya | Transporter | 3ORG<sup>12</sup> | Yes |
| bCLC-K | E1B792 | Eukarya | Channel | 5TQQ<sup>18</sup>, 5TR1<sup>18</sup> | Yes |
| hCLC-1 | P35523 | Eukarya | Channel | 6COY<sup>20</sup>, 6QV6<sup>22</sup>, 6QVB<sup>22</sup>, 6QVC<sup>22</sup>, 6QVD<sup>22</sup>, 6QVU<sup>22</sup> | No |
| hCLC-2 | P51788 | Eukarya | Channel | 7XF5<sup>29</sup>, 7XJA<sup>29</sup>, 8TA2<sup>31</sup>, 8TA3<sup>31</sup>, 8TA4<sup>31</sup>, 8TA5<sup>31</sup>, 8GQU<sup>31</sup> | No |
| mCLC-3 | P51791 | Eukarya | Transporter | 8JEV<sup>34</sup>, 8JGK<sup>34</sup>, 8JGJ<sup>34</sup>, 8JGL<sup>34</sup>, 8JGS<sup>34</sup>, 8JGV<sup>34</sup> | No |
| hCLC-6 | P51797 | Eukarya | Transporter | 8JPO<sup>35</sup>, 8JPJ<sup>35</sup>, 8JPR<sup>35</sup> | No |
| hCLC-7 | P51798 | Eukarya | Transporter | 7BXU<sup>24</sup>, 7CQ5<sup>25</sup>, 7CQ6<sup>25</sup>, 7CQ7<sup>25</sup>, 7JM7<sup>27</sup>, 8HVT<sup>32</sup> | No |
| atCLC-a | P92941 | Eukarya | Transporter | 7XA9<sup>30</sup>, 8IAB<sup>33</sup>, 8IAD<sup>33</sup> | No |
| ggCLC-7 | Q5ZL60 | Eukarya | Transporter | 7JM6<sup>27</sup> | No |

**Supplementary Table 2.** List of alignment sites used in distance matrix analysis and the corresponding CLC-ec1 residues.

| Structural Element | CLC-ec1 Residues | Alignment Sites |
| --- | --- | --- |
| Helix αB | 34-66 | 1-33 |
| Helix αC | 76-99 | 34-57 |
| αC-D linker | 100-109 | 58-67 |
| Helix αD | 110-117 | 68-75 |
| Helix αE | 125-141 | 76-92 |
| Helix αF | 148-166 | 93-111 |
| Helix αG | 172-189 | 112-129 |
| Helix αH | 194-203 | 130-139 |
| Helix αI | 216-232 | 140-156 |
| Helix αJ | 253-281 | 157-185 |
| Helix αK | 290-303 | 186-199 |
| Helix αL | 318-324 | 200-206 |
| Helix αM | 331-349 | 207-225 |
| Helix αN | 357-377 | 226-246 |
| Helix αO | 387-402 | 247-262 |
| Helix αP | 407-416 | 263-272 |
| Helix αQ | 423-437 | 273-287 |
| Helix αR | 444-450 | 288-294 |

**Supplementary Table 3.**
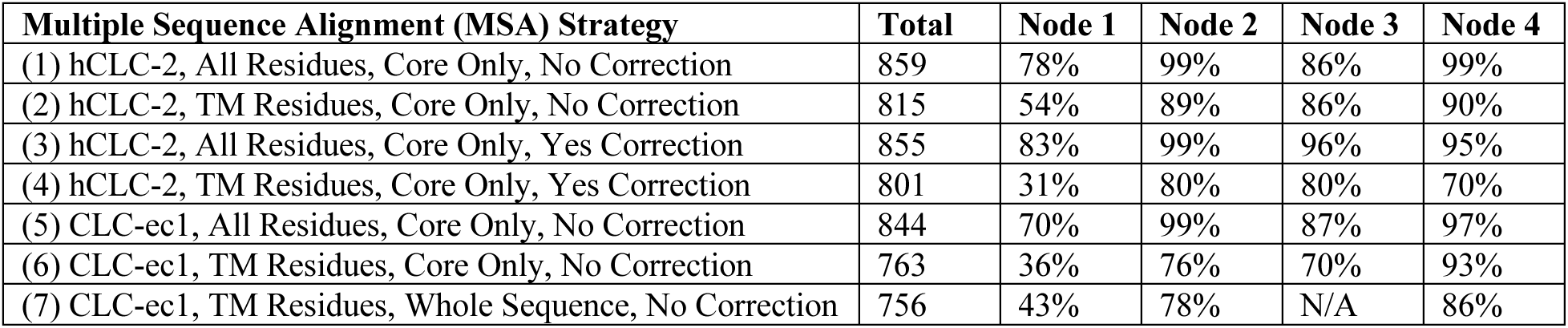
Sum of node support values for different multiple sequence alignment (MSA) methods.

